# A snapshot of progenitor-derivative speciation in action in *Iberodes* (Boraginaceae)

**DOI:** 10.1101/823641

**Authors:** Ana Otero, Pablo Vargas, Virginia Valcárcel, Mario Fernández-Mazuecos, Pedro Jiménez-Mejías, Andrew L. Hipp

## Abstract

- Traditional classification of speciation modes has focused on physical barriers to gene flow. While allopatry has been viewed as the most common mechanism of speciation, parapatry and sympatry, both entail speciation in the face of ongoing gene flow and thus both are far more difficult to detect and demonstrate. *Iberodes* (Boraginaceae, NW Europe) with a small number of recently derived species (five) and contrasting morphological traits, habitats and distribution patterns constitutes an ideal system in which to study drivers of lineage divergence and differentiation.
- To reconstruct the evolutionary history of the genus, we undertook an integrative study entailing: (i) phylogenomics based on restriction-site associated DNA sequencing (RAD-seq), (ii) morphometrics, and (iii) climatic niche modelling.
- Key results revealed a history of repeated progenitor-derivative speciation, manifesting in paraphyletic pattern within *Iberodes*. Climatic niche analyses, together with the morphometric data and species distributions, suggest that ecological and geographical differentiation have interacted to shape the diversity of allopatric and parapatric distributions observed in *Iberodes*.
- Our integrative study has enabled to overcome previous barriers to understanding parapatric speciation by demonstrating the recurrence of progenitor-derivative speciation in plants with gene flow and ecological differentiation, explaining observed parapatry and paraphyly.

## INTRODUCTION

Understanding what modes of speciation dominate diverse clades across the tree of life has been a contentious domain of evolutionary biology for decades (e.g. Templeton, 1981; Rieseberg & Brouillet, 1994; Butlin *et al.*, 2008; Horandl & Stuessy, 2010). Reproductive barriers are the most frequently used criteria for distinguishing among three overarching modes of speciation (Futuyma, 2009). Speciation of adjacent populations in the absence of clear physical or reproductive barriers is frequently explained under the process of (1) parapatric speciation, which entails the evolution of reproductive isolation in the face of limited but ongoing gene flow (Coyne & Orr, 2004, p. 112). Parapatric speciation can be considered intermediate between allopatric and sympatric speciation. In (2) allopatric speciation, gene flow is prevented by geographic barriers between populations. In (3) sympatric speciation, partial or complete reproductive isolation arises between subsets of a single population with overlapping ranges, without spatial segregation. Sympatric speciation often entails significant levels of gene flow at the early stages of differentiation (Gavrilets, 2003; Futuyma, 2009). Whereas allopatry is widely viewed to be the most common mode of speciation (Coyne & Orr, 2004; Vargas *et al.*, 2018), the relative importance of parapatry and sympatry has been the subject of considerable debate (Fitzpatrick *et al.*, 2008; Fitzpatrick *et al.*, 2009; Arnold, 2015) in large part due to the difficulty of (1) resolving phylogenetic relationships among very closely related populations that may still be exchanging genes at a low level, and (2) quantifying the amount of gene flow between populations during population divergence (Barluenga *et al.*, 2006; Fontdevilla, 2014). In the last decade, new genomic, phylogenetic, and ecological tools have made non-allopatric modes of speciation increasingly amenable to study, allowing us to explore with statistical rigor a wider range of speciation scenarios (e.g. Savolainen *et al.*, 2006; Chozas *et al.*, 2017; Zheng *et al.*, 2017). In particular, there are some critical points of evidence that together provide support for a parapatric speciation scenario: (1) phylogenetic relationships usually reflect a progenitor-derivative (budding) pattern in which the more widely distributed species is the living progenitor of the more restricted one; (2) the absence of any physical barrier to gene-flow in both present and past times, i.e. distribution ranges are adjacent; (3) interspecific genetic differentiation that putatively discards processes of secondary contact; (4) alternative forces of reproductive isolation factors—e.g., differential climatic niches, reproductive exclusion, ploidy variation, among others—are present.

Ecological differentiation is often associated with non-allopatric speciation (Sobel *et al.*, 2010; Nosil, 2012; Stankowsky *et al.*, 2015), particularly in plants, because a sessile habit makes plants more sensitive to fine-scale environmental heterogeneity (Anacker & Strauss, 2014). Both the influence of niche conservatism (i.e. retention of ancestral ecological characteristics of species over time, Peterson *et al.*, 1999) and niche evolution (i.e. adaptation of lineages to changes in the environment, Donoghue & Edwards, 2014) have been argued to play a role in speciation and lineage diversification (Wiens & Graham, 2005; Cavender-Bares, 2019). Nevertheless, the contribution of ecological factors as primary drivers of speciation remains more elusive (Mayr, 1954; Pyron & Burbrink, 2010; Yin *et al.*, 2016).

The advent of inexpensive phylogenomic approaches in the last decade have made the genetic side of testing recent speciation scenarios tractable (McCormack *et al.*, 2013; McVay *et al.*, 2017a). High Throughput Sequencing (HTS) approaches make it possible to analyze loci sampled from the entire genome in reconstructing species trees (Fernández-Mazuecos *et al.*, 2018) and the history of speciation in the face of gene flow (Leroy *et al.*, 2017; Folk *et al.*, 2018; Crowl *et al.*, 2019). Restriction-site associated DNA sequencing (RAD-seq) has contributed particularly strongly to our understanding of recent evolutionary processes, especially in non-model organisms, since a reference genome is not needed for accurate phylogenetic inference (e.g. Fitz-Gibbon *et al.*, 2017). RAD-seq has as a consequence been useful in inferring complex speciation histories: repeated cycles of island connectivity and isolation (the pump hypothesis; Papadopoulou & Knowles, 2015), incipient sympatric speciation (Kautt *et al.*, 2016), allopatric speciation despite historical gene flow (Maguilla *et al.*, 2017), ancient introgression among now-extinct species (McVay *et al.*, 2017b).

The Mediterranean subendemic genus *Iberodes* M.Serrano, R.Carbajal & S.Ortiz (Boraginaceae, Cynoglossoideae; Serrano *et al.*, 2016) provides an excellent system for studying recent speciation, morphological and ecological differentiation because it is clearly demonstrated to be a recently-derived genus (Chacón *et al.*, 2017; Otero *et al.*, 2019a). *Iberodes* comprises five species and one subspecies that differ in geographic ranges (including allopatric and parapatric species) and habitat (ranging from forest understories to scrublands and coastal dunes). Moreover, all species are endangered and narrowly endemic, except for the widely-distributed *I. linifolia* of the Iberian Peninsula and southeastern France (Talavera *et al.*, 2012). Sanger sequencing has failed to resolve phylogenetic relationships among *Iberodes* species because the markers used to date provide minimal variability (Otero *et al.*, 2014; Holstein *et al.*, 2016a; Holstein *et al.*, 2016b; Otero *et al.*, 2019a). The question consequently remains as to what is the relative importance of extrinsic geographical and ecological barriers to the complex speciation patterns observed in *Iberodes*. Given the narrow endemicity in most *Iberodes* species, we hypothesize that geographic barriers rather than ecological factors have been predominant in the speciation history of *Iberodes*. To test this hypothesis, we aim to: (1) reconstruct phylogenetic relationships of the five species; (2) infer speciation patterns and lineage diversification during the last geological epochs comparing diverse morphologies and ecologies; and (3) evaluate the relative contribution of geographical and ecological processes in speciation.

## MATERIAL AND METHODS

### Taxon sampling

Based on species distributions, between one and four individuals from one to nine populations were sampled for each of the five species of *Iberodes* including ssp. *gallaecica* and ssp. *littoralis* of *I. littoralis*) (Fig. 1, Table S1, in Supporting Information). Samples of *I. littoralis*, *I. kuzinskyana* and *I. brassicifolia* were collected in the field during 2015 and stored in silica gel until extraction. Since these species are considered threatened according to IUCN criteria (Lopes & Carvalho, 1990; Serrano & Carbajal, 2003; ICNB, 2007; Moreno, 2008; IUCN, 2019; INPN, 2019), permits were obtained prior to collecting. Mature seeds of *I. commutata* were collected in 2015 and germinated in a glasshouse to obtain green leaves for DNA extraction. Both field and herbarium materials were used to represent the large distribution of *I. linifolia* (Table S1). Three species (17 individuals) of the tribe Omphalodeae were included as the outgroup (Otero *et al.*, 2019b): *Omphalodes nitida*, *Myosotidium hortensia*, and *Gyrocaryum oppositifolium* (Table S1).

**Fig. 1.**
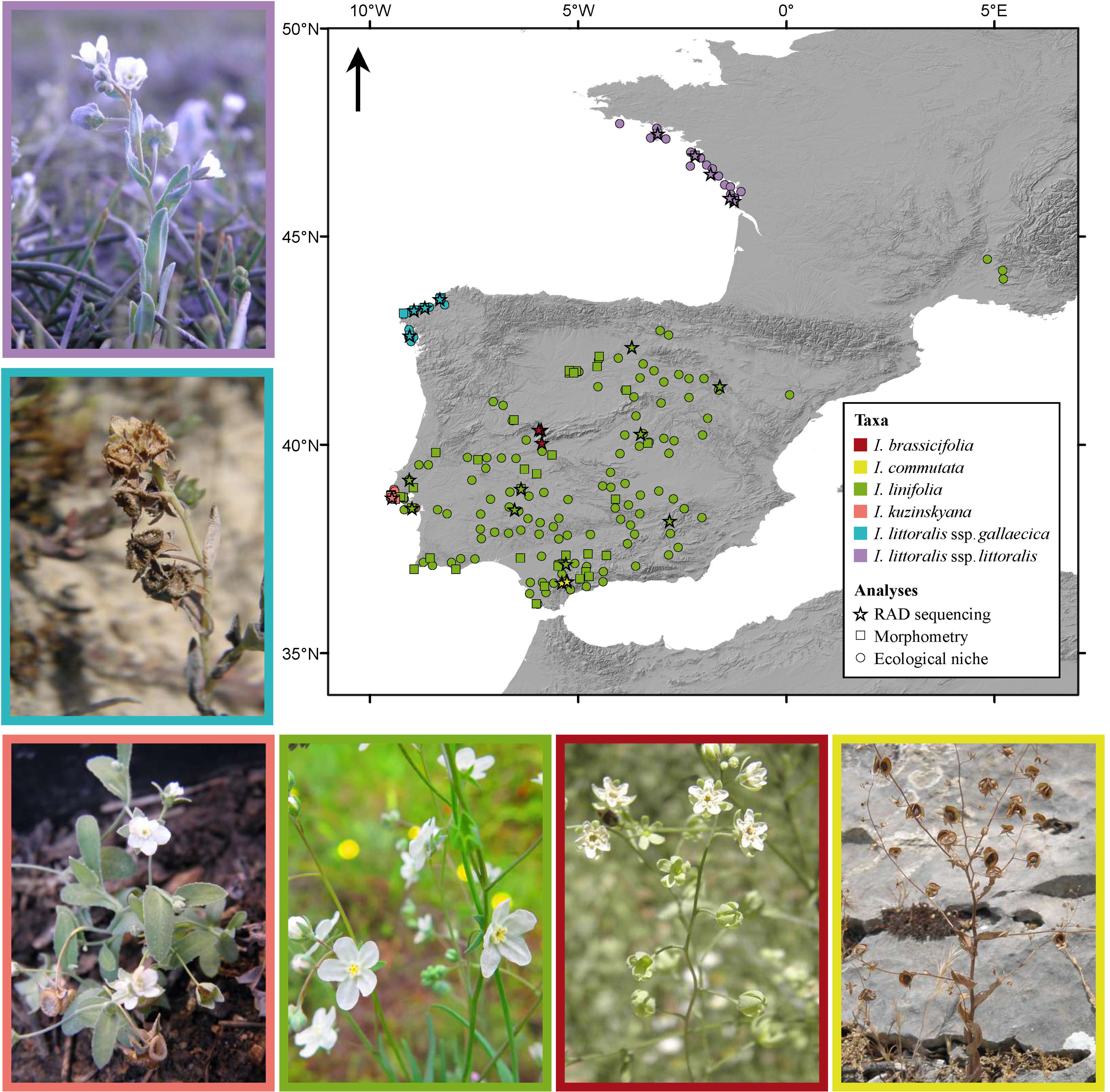
Locations of the *Iberodes* species sampled for the different analyses performed. The star symbols indicate locations for RAD-sequencing, squares represent the locations for the morphometric analysis, and circles indicate locations for the ecological niche modelling. Color legend for each taxa of *Iberodes* is shown in the figure. Images of the six taxa are framed following each taxa color in the map.

### DNA extraction and RAD library preparation

DNA was extracted from leaf tissue using a modified CTAB protocol from Doyle and Doyle (1987) and Shepherd and Mc Lay (2011), including a precipitation step in isopropanol and ammonium acetate 7.5M at −20°C overnight to improve yield. DNA extractions were visualized and quantified with NanoDrop 2000/2000c (Thermo-Fisher, Waltham, Massachusetts) and Qubit Fluorometric Quantification (Thermo-Fisher, Waltham, Massachusetts) in the *Instituto de Investigaciones Biomédicas* (IIBm, CSIC-UAM). Final DNA sample concentrations were standardized to 10 ng/ul. Preparation of single-end RAD-seq libraries using restriction enzyme *Pst*I from genomic DNA was conducted at Floragenex Inc. (Eugene, Oregon) following Baird *et al.* (2008) and sequenced as single-end, 100-bp reactions on an Illumina HiSeq 2500 at the University of Oregon Genomics & Cells Characterization Core Facility (library C606). Processed data were returned in the Illumina 1.3 variant of the FASTQ format, with Phred quality scores for all bases.

### Data clustering

Sequences from Illumina were analyzed in *PyRAD* v. 3.0.6, a *de novo* clustering pipeline with seven main steps that filter and cluster putatively orthologous loci from RAD sequencing reads (Eaton, 2014). This pipeline is able to cluster highly divergent sequences, taking into account indel variation and nucleotide (SNP) variation. It is thus well-suited to the inter- and intraspecific scales addressed in our study. As no evidence of polyploidy was found from flow cytometry, we did not assess sensitivity of paralog detection to sequencing depth or number of heterozygotic positions allowed per loci or shared among individuals. Base calls with a Phred Quality score <20 were replaced by the ambiguous base code (N). Three similarity percentages were set for the clustering (c) threshold of reads: 85%, 90% and 95%. Four levels of minimum coverage (m) of a sample in a final locus (4, 12, 20, 28) were tested for each of the three percentages of clustering similarity, resulting in a fully factorial set of 12 matrices from the combination of filtering parameters, so we could assess sensitivity of our phylogenetic inferences to clustering assumptions. Consensus base calls were made for clusters with a minimum depth of coverage >5. From each parameter combination we obtained three kinds of output: (1) a matrix with all loci retrieved for each individual (concatenated matrix), (2) a matrix with single nucleotide polymorphisms (SNP matrix), and (3) a matrix with one SNP per locus (unlinked SNP matrix). The loci matrix was explored with the RADami (Hipp, 2014) and vegan (Oksanen *et al.*, 2013) packages of R version 3.3.2 (R Core Team, 2013).

### Phylogenetic analysis

Maximum likelihood analyses of concatenated matrices obtained in *PyRAD* were performed in RAxML v8.2.10 (Stamatakis, 2014) through the Cipres Portal (Miller *et al.*, 2010). We used rapid bootstrapping, the GTRGAMMA model, and automatic-stop bootstrapping with the majority rule criterion. We conducted a maximum likelihood analysis for each of the 12 *PyRAD* concatenated matrices. Finally, we compared the topologies based on average bootstrap support (BS). Between the two topologies with average BS>98%, we chose the one maximizing the number of loci (HBS tree) for the remaining phylogenetic-based analyses. Moreover, we used the unlinked SNP matrix from the HBS tree (HBS matrix) to perform a coalescent-based phylogeny using SVDquartets (Chifman & Kubatko, 2014) in PAUP (Swofford, 2001) through the Cipres portal (Miller *et al.*, 2010). For SVDquartets we grouped individuals based on main clades obtained from concatenated analysis in RAxML and evaluated all possible quartets with 100 bootstrap replicates.

### Estimates of divergence times

We used penalized likelihood as implemented in TreePL (Smith & O’Meara, 2012) to estimate a time-calibrated tree for all bootstrap trees (with branch lengths) obtained in RAxML from the HBS tree. TreePL is suitable for divergence time estimation when dealing with large amounts of data, such as those yielded by high throughput sequencing RAD-sequencing (Zheng & Wiens, 2015). Two calibration points were used: (1) tribe Omphalodeae (root) (minimum age (minAge) = 9.062; maximum age (maxAge) = 24.4097) and (2) clade *Iberodes* (ingroup) (minAge = 0.6591; maxAge = 3.7132) based on the averaged ages and standard deviation inferred in Otero *et al.* (2019a). While secondary calibrations tend to underestimate the uncertainty of the age estimates in the trees from which they are derived, use of multiple secondary calibrations with the full range of uncertainty found in the original study have the potential to recover more accurate clade ages (Schenk, 2016). Average node ages over all posterior trees from the latter study were calculated using *treeStat* from the BEAST v 1.8.2 package (Drummond *et al.*, 2012). We first conducted an analysis under the “prime” option to select the optimal set of parameter values, using random subsample and replicate cross-validation (RSRCV) to identify the best value for the smoothing parameter, lambda. For this first analysis we used the thorough analysis option and set 200,000 iterations for penalized likelihood and 5,000 iterations for cross validation. We then repeated the same thorough analysis and iterations but setting the smoothing value of lambda to 10, gradient based (opt) and auto-differentiation based optimizers (optad) to five, and auto-differentiation cross-validation-based optimizers to two (optcvad). Although TreePL does not draw a prior distribution for the phylogenetic tree, we took into account the bootstrapped variance in topology and branch length by running the same TreePL analysis for each bootstrap tree and then a maximum clade credibility tree using the mean heights was reconstructed in *TreeAnnotator* from the BEAST v. 1.8.2 package (Drummond *et al.*, 2012). Our dating analysis thus accounts for the uncertainty in branch length that is due to variance in molecular substitution process across the RAD-seq loci used.

### Species-level analyses

Phylogenetic inferences showed two cases of paraphyly within the lineage of *I. linifolia* that involved three other taxa (*I. kuzinskyana*, *I. littoralis* ssp. *littoralis* and ssp. *gallaecica*; see results). To evaluate these cases of paraphyly, a set of analyses were done to analyze the degree of genetic, morphological and ecological differentiation among the four taxa involved. Therefore, the following analyses were performed on a subset of data including only these four taxa.

#### Genetic structure

The HBS matrix was used to perform a Discriminant Analysis of Principal Components (DAPC) to identify and describe clusters of genetically related individuals (Jombart *et al.*, 2010). The format of the HBS matrix was converted from *vcf* to *genlight* with vcfR (Knaus & Grünwald, 2017). DAPC analysis was performed in adegenet (Jombart, 2008). We fixed the number of clusters to four, based on the natural groups of four taxonomic species and subspecies (the Linifolia clade, see results). We used the function ‘optim.a.score’ to estimate the optimal number of principal components for discriminant analysis. We represented genetic differentiation using scatterplots of the principal components and a barplot representing assignment probabilities of individuals to groups (compoplot). In addition, analysis of molecular variance (AMOVA) was performed in Arlequin v. 3.5.2.1 (Excoffier & Lischer, 2010). AMOVA was run to evaluate the level of genetic differentiation of the paraphyletic *I. linifolia* as a whole with respect to the other two species (*I. kunzinskyana* and *I. littoralis*) of the Linifolia clade. Therefore, we considered only two groups, *I. linifolia* on one hand, and the two coastal species together (*I. kusinskyana* and *I. littoralis*) on the other. Finally, to evaluate a potential role of historical introgression in determining paraphyly, we conducted D-statistic tests in PyRAD (see Methods S1 for details).

#### Morphometric evaluation

We performed a morphometric exploration using principal component analysis (PCA). We consulted the taxonomic literature to identify diagnostic characters in *Iberodes* (Talavera *et al.*, 2012). We measured 12 diagnostic macromorphological vegetative and reproductive characters for 111 individuals from 58 vouchers (two different individuals per voucher except for 5 vouchers with only one individual). We averaged measures from each two individuals of the same voucher. As a result, we obtained a final matrix of 58 accessions from 47 locations (74 individuals from 35 locations of *I. linifolia*, 14 from five locations of *I. kuzinskyana*, and 23 from seven locations of *I. littoralis* (including the two subsppecies) (Fig. 1, Table S2). Ten morphological characters were quantitatively continuous: four vegetative characters (length of stem, leaves and pedicels, and width of leaves) and six reproductive characters (length of fruiting calix, width and length of nutlet, length of nutlet margin, length of the nutlet margin teeth, and hair length). In addition, one binomial character (presence or absence of bracts) and one quantitative, discrete character (hairs per mm^2^ on the abaxial surface of the nutlet) were also considered.

PCA was run using the function ‘prcomp’ from Stats R package (R Core Team, 2013). Results were visualized with the function ‘ggbiplot’ in the ggbiplot R package (Vu, 2011).

#### Climatic niche differentiation and distribution modelling

Occurrences of the four taxa were collected from GBIF (https://www.gbif.org/). We filtered the resulting dataset by removing points suspected to be incorrect, such as those placed clearly outside of the known species distribution range (based on taxonomic literature and databases; Fernández & Talavera, 2012; Muséum National d’Histoire Naturelle, 2003-2019). Additional geographic coordinates were obtained from herbarium material and our own fieldwork data. We also estimated the geographic coordinates of non-georeferenced herbarium specimens, whenever the locality provided could be pinpointed with an accuracy of ± 5 km (Table S3, Fig. 1). To reduce sampling bias, we filtered the dataset by randomly removing points that were within 0.2° latitude or longitude of each other for *I. linifolia* and 0.1° for *I. littoralis* ssp. *littoralis* and ssp. *gallaecica*. Filtering was not applied for *I. kuzinskyana* because of the low number of occurrences. The final dataset after filtering included 115 occurrences of *I. linifolia*, 11 of *I. kuzinskyana*, 16 of *I. littoralis* ssp. *littoralis* and 11 of ssp. *gallaecica*.

We obtained 19 bioclimatic variables (resolution of 30”) from WorldClim 1.4 (http://www.worldclim.org/; Hijmans *et al.*, 2006) for the study region (34° to 50° N; 11° W to 7° E). To avoid collinearity among bioclimatic variables, we first excluded variables displaying a high correlation coefficient with other variables (|r| > 0.7) in the study area (Dormann *et al.*, 2013). Then, we calculated the variance inflation factor (VIF) for each remaining variable using the HH package in R (Heiberger, 2017) and selected variables with VIF <5 following Benítez-Benítez *et al.* (2018). As a result, the following six variables were selected: bio1, bio3, bio4, bio8, bio12 and bio14. We assessed climatic niche overlap among the four taxa by pairwise comparisons. The E-space (i.e. environmental space resulting from the six climatic variables) of each taxon was analyzed in a Principal Component Analysis (PCA) as implemented in the R package ecospat (Di Cola *et al.*, 2017). To visualize niche differences along each of the first two principal components, the result was plotted using the niceOverPlot function (Fernández-López & Villa-Machío, 2017), which relies on the ggplot2 package (Wickham, 2016). We calculated values of Schoener’s index (D) as a measure of niche overlap (Schoener, 1968; Warren *et al.*, 2008). Then, we conducted tests of niche equivalency and niche similarity using ecospat (Di Cola *et al.*, 2017). The niche equivalency test evaluates whether the observed niche overlap is significantly different from a null simulated by randomly reallocating the occurrences of both entities between their ranges (Broennimann *et al.*, 2012; Warren *et al.*, 2008). The niche similarity test checks whether the overlap between two niches is significantly different from the overlap obtained if random shifts within each environmental space are allowed (Schoener, 1968; Warren *et al.*, 2008). In both cases, we tested for niche divergence by using the alternative=“lower” option. All tests were based on 100 iterations. In similarity tests, both ranges were randomly shifted (rand.type=1).

In addition, we modeled the potential range of the four taxa of the Linifolia clade under present conditions, which was then projected to last interglacial (LIG, c. 120–140 kya) and last glacial maximum (LGM, c. 21 kya) conditions. To this end, we used the same six WorldClim bioclimatic variables employed for niche differentiation analyses, including present layers, LIG layers from Otto-Bliesner *et al.* (2006) and LGM layers based on the CMIP5 project (three global climate models (GCMs): CCSM4, MIROC-ESM and MPI-ESM-P). Resolution was 30“ for current and LIG, and 2.5’ for LGM layers. We performed species distribution modeling (SDM) using the maximum entropy algorithm, as implemented in Maxent v3.4 (Phillips *et al.*, 2006). We used the same occurrences for each taxon employed for niche differentiation analyses (see above). 80% of occurrences for each taxon were used for model training and 20% for model evaluation. For each taxon, ten subsample replicates were run by randomly partitioning the data into a training set and evaluation set, and a mean model was calculated. For the LGM, an average of the three GCMs was calculated. Logistic outputs were converted into presence/absence using the maximum training sensitivity plus specificity logistic threshold.

#### Ploidy level estimation

Flow cytometry was used to estimate genome size, explore the possible role of polyploidization in speciation, and assess the risk of clustering paralogs in RAD-seq *de novo* clustering. Earlier cytogenetic studies for three *Iberodes* species showed them to be diploids (2n=28 for *I. linifolia*, *I. commutata* and *I. kuzinskyana* (Franco, 1984; Saly, 1997; Talavera *et al.*, 2012), and aneuploidy was inferred for *I. littoralis* based on its chromosome count of 2n=24 (Fernández-Casas, 1975). Nevertheless, some of these references do not cite the plant sources, making it difficult to assess the connection of cytotypes to populations. Individuals of the five *Iberodes* species were cultivated at the glasshouse and five individuals per species were sampled. In addition, samples of the diploid *Solanum lycopersicum* of known genome size (2n=24, 2C=1.88-2.07 pg, Grandillo *et al.*, 2011) were also cultivated to compare standardized results with our study species and estimate genome sizes (Doležel & Bartoš, 2005). We performed a two-step nuclear DNA Content Analysis using Partec Buffer (de Laat *et al.*, 1987) and DAPI fluorochrome through a Cell Lab Quanta SC flow cytometer (Beckman Coulter, Fullerton, CA, USA) equipped with a mercury lamp following the protocol of Doležel *et al.* (2007). Samples were visualized through the Cell Lab Quanta SC Software package. FL1 detector (530 nm / 28 nm; DAPI emission maximum=461) and FL1-Area was used as directly correlated to DNA content. A minimum number of 1000 nuclei for G1 peak were analyzed. Only histograms with a coefficient of variation (CV) lower than 10% for the G1 peak were accepted.

## RESULTS

### Phylogenetic analysis

After filtering and processing all samples under the different parameter combinations in *PyRAD*, 12 matrices of 54 taxa each but varying in numbers of loci and SNPs were analyzed (see Table S4). The same topology was obtained in RAxML from each of the 12 matrices, differing only in bootstrap support (BS) values for the nodes (Fig. S1). The highest BS supports and highest number of loci were obtained for the c95m20 matrix (95% similarity within clusters and minimum coverage of 20 samples for a final locus). Consequently, this matrix was used for the remaining phylogenetic analyses (HBS matrix). *Iberodes brassicifolia* was inferred to be the earliest-diverging lineage, sister to a clade that contains *I. commutata* sister to another subclade with the remaining species (Fig. 2). Paraphyly was inferred for *I. linifolia* since a clade including *I. kuzinskyana* and *I. littoralis* is nested within. Likewise, paraphyly was inferred for *I. littoralis* ssp. *gallaecica* since ssp. *littoralis* is nested within populations of subsp. *gallaecica* (Fig. 2).

Taxon groups for coalescent-based phylogenetic inference (SVDquartets) were based on the eight main lineages recovered using RAxML (*Iberodes brassicifolia*, *I. commutata*, populations of *I. linifolia* from Seville, core clade of *I. linifolia*, populations of *I. linifolia* from southern Spain, populations of *I. linifolia* from central Portugal, *I. kusinskyana*, and *I. littoralis*, Fig. S2). The species tree topology was mostly congruent with the concatenated phylogenetic topology. The only difference was the placement of *I. linifolia* populations from Seville and southern Spain, which were nested within the core of *I. linifolia* (Fig. S2). In any case, paraphyly of *I. linifolia* is still inferred because of the nested position of *I. kuzinskyana* and *I. littoralis* sister to populations of *I. linifolia* from Portugal (Fig. S2).

**Fig. 2.**
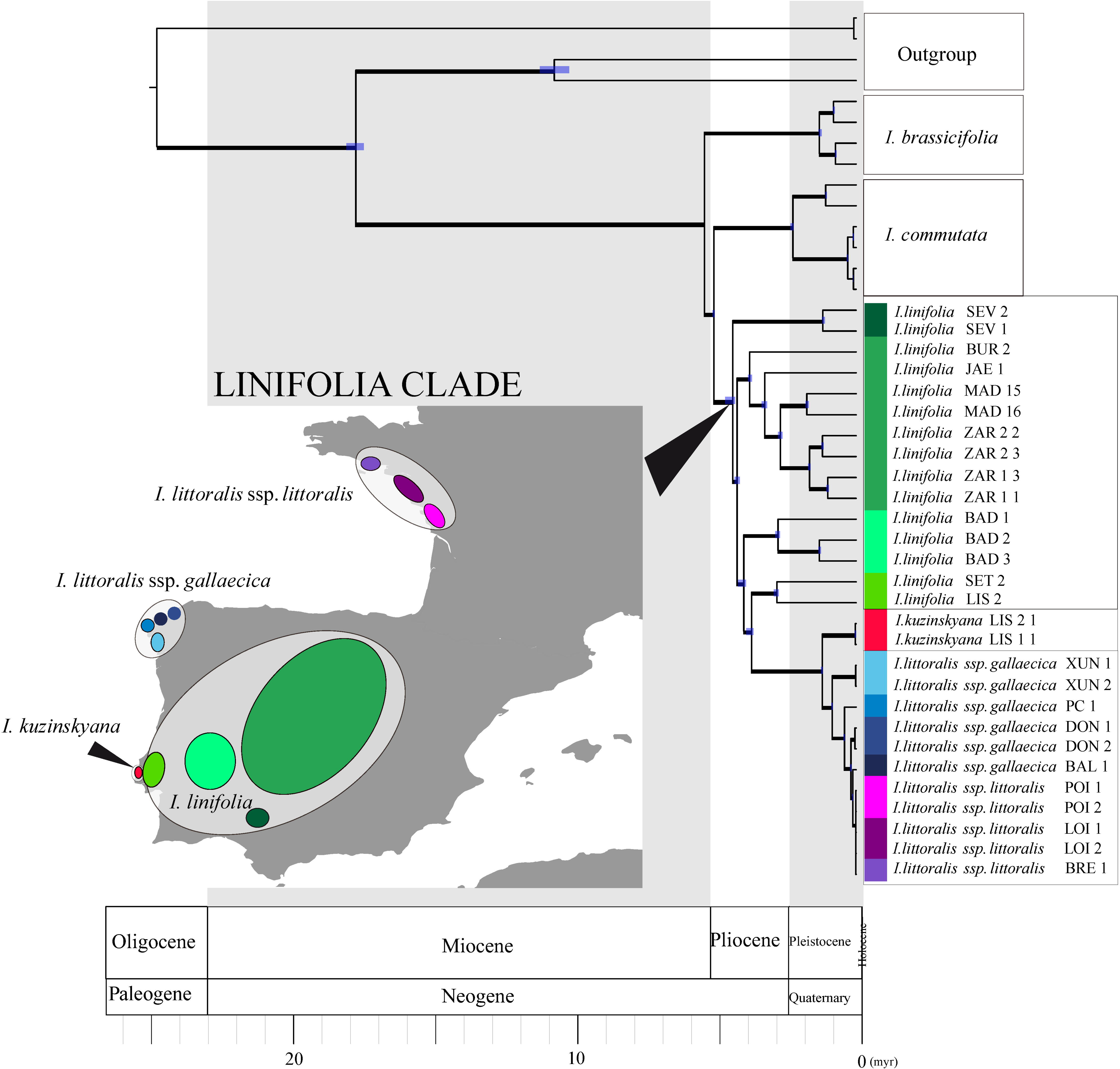
Time-calibrated phylogeny obtained from TreePL by using maximum clade credibility from 1000 bootstrap trees of the maximum likelihood analysis of c95m20 matrix. Branch thickness indicates the ML bootstrap supports above 98. Node bars in blue indicate the node age ranges taking into account the branch length variance along the 1000 bootstrap trees. No branch length variation was inferred for nodes without blue bar. The international stratigraphic scale is included from 26 myr until present. Text codes for each individual location are detailed in the Table S1. The color for each lineage of the Linifolia clade is geographycally represented through the colored ellipses of the map. Grey ellipses group the different lineages of the four taxa within the Linifolia clade.

### Divergence time estimates

The stem age estimated for *Iberodes* was 17.56 myr (17.26-17.86 Bootstrapped Variance, BV) with a crown age of 5.32 myr (5.32-5.32 BV) (Fig. 2). Crown age estimates for *I. brassicifolia* and *I. commutata* were 1.29 (1.21-1.33 BV) and 2.22 myr (2.17-2.28 BV), respectively. The crown age of the Linifolia clade (*I. linifolia* s.l., *I. kuzinskyana* and *I. littoralis*) was inferred to be 4.33 myr (4.25-4.59 BV). Divergence time between Portuguese populations of *I. linifolia* and ancestor of the coastal species (*I. kuzinskyana* and *I. littoralis*) was estimated at 3.67 myr (3.60-3.85 HPD). The origin of *I. littoralis* ssp. *littoralis*, possibly from an *I. littoralis* ssp. *gallaecica* ancestor, was inferred to have taken place ca. 10,000 years ago (Fig. 2).

### Genetic structure

We set four as the *a priori* number of clusters (K=4) based on the four taxa circumscribed within the Linifolia clade. DAPC analysis identified only one optimal principal component for the scatterplot which revealed two well-differentiated clusters: *I. linifolia* (LIN) on the one side, and the coastal *I. kuzinskiana* (KUZ) and *I. littoralis* (LIT_LIT, LIT_GAL) on the other (Fig. 3c). Within the coastal species cluster, *I. kuzinskyana* occupied a position closer to *I. linifolia*, while the two subspecies of *I. littoralis* were intermingled (Fig. 3c). Compoplot assigned all individuals of *I. linifolia* to a separate cluster, without admixture (Fig. 3c). The other three clusters were shared among coastal species with different membership probability (Fig. 3c). One of the two individuals of *I. kuzinskyana* was uniquely assigned to one of the clusters, whereas the other individual showed a mixed assignment, with admixture of the three genetic groups (Fig. 3c). Most individuals (nine) of both subspecies of *I. littoralis* were placed in the same two clusters, and two of them showed admixture with the same cluster shared by the two samples of *I. kuzinskyana* (Fig. 3c). Given the interdigitated pattern of the two subspecies of *I. littoralis*, we repeated the analysis by considering the two subspecies as part of a single group (K=3) to evaluate the genetic clustering of *I. littoralis* as a genetic group. Compoplot for K=3 assigned most individuals (nine) of *I. littoralis* to the same cluster, while two samples had mixture assignment probability to the cluster of *I. kuzinskyana* (Fig. S3). Likewise, AMOVA of the two groups (*I. linifolia* vs coastal species) showed significant among-group variance (58.88%) higher than within-population variance (38.09%) or variance within populations among groups (3.03%), with a fixation index of F_ST_= 0.619 (Table 1). D-statistic tests did not conclusively support a role of historical introgression in determining paraphyletic pattern since ancient introgression was inferred indiscriminately among lineages of *I. linifolia* and both coastland species (see Methods S1).

**Table 1.** Analysis of molecular variance (AMOVA) results for individuals from two contrasted groups: (1) mainland, *I. linifolia* and (2) coastal, *I. kuzinskyana* and *I. littoralis.*

| Source of Varitation | d.f. | Sum of squares | Variance components | Percentage of variation |
| --- | --- | --- | --- | --- |
| Among groups | 1 | 1602.666 | 106.33509 Va | 58.88 |
| Among populations within groups | 2 | 181.372 | 5.47321 Vb | 3.03 |
| Within populations | 24 | 1651.033 | 68.79306 Vc | 38.09 |
| Total | 27 | 3435.071 | 180.60135 |  |

Note: Variance components are as in Excoffier *et al.* (1992). Fst = 0.61909; Fsc = 0.07370; Fct = 0.58878.

**Fig. 3.**
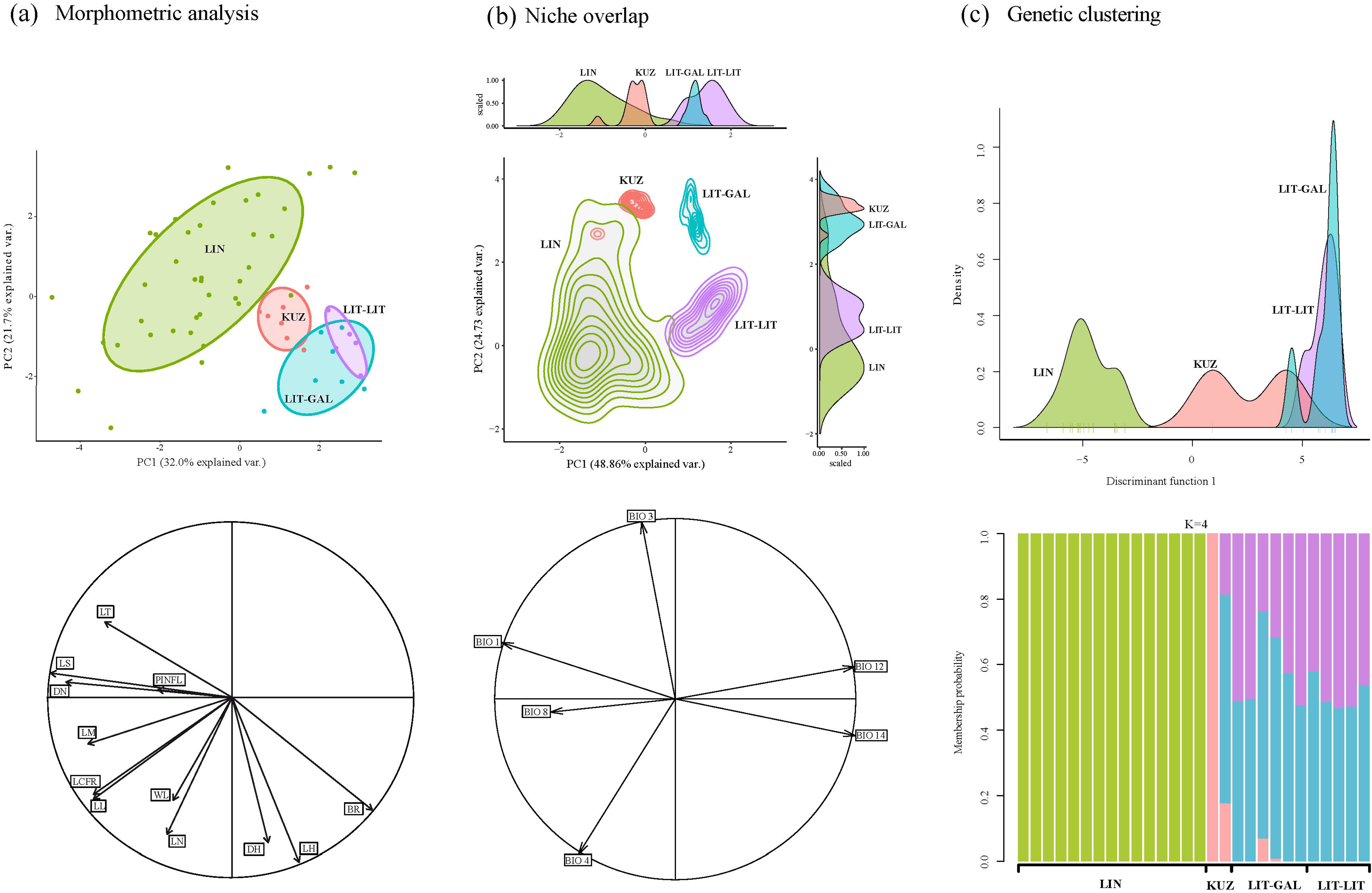
Morphometric, niche overlap and genetic clustering analyses of the Linifolia clade. Codes for each taxa are: LIN for *I. linifolia*, KUZ for *I. kuzinskyana*, LIT-GAL for *I. littoralis* ssp. *gallaecica*, and LIT-LIT for *I. littoralis* ssp. *littoralis*. (a) Morphometric analysis. Principal Component Analaysis (PCA) of the 12 morphologic chartacters: LS, length of the stem; LL, length of the leaf; WL, wide of the leaf; PINFL, pedicel of inflorescence; LCFR, length of fruit calix; DN, diameter of the nutlet; LN, length of the nutlet; LM, length of the margin nutlet; LT, length of the margin teeth; DH, density of trichomes; LH, length of trichome; BR: absence=0, presence=1 of flowering bracts). The contribution of each trait to the two principal components (PC1, PC2) as well as the positive or negative sense of the relationship is represented through the length of each arrow and the directionality of the arrow, respectively. The percentages of the variability explained by the two PCs are indicated close to the axes. (b) Niche overlap analysis. PCA of the six climatic variables obtained from WorldClim. BIO1, Annual Mean Temperature; BIO3, Isothermality; BIO4, Temperature Seasonality; BIO8, Mean Temperature of Wettest Quarter; BIO12, Annual Precipitation; BIO14, Precipitation of Driest Month. The strength of contribution of each variable to PCs and the sense of the relationship is represented, as explained above, through the length and the directionality of the arrows. The percentages of the variability explained by the two PCs are indicated close to the axes. Values of the variables are also represented for each PC separately following Fernández-López and Villa-Machío (2017). (c) Genetic clustering. Scatterplot from the DAPC analysis showing the genetic differentiation retained in one PC as the optimal number of PC for the discriminant function. Compoplot for K=4 is also represented indicating membership probability to each genetic cluster for each taxa.

### Morphometric analysis

Principal Component Analysis (PCA) revealed morphological differentiation between the three taxa. *Iberodes linifolia* exhibited the greatest morphological variation, ranging from the two extremes of both PC1 and PC2 compared to the more restricted variation of coastal species that is concentrated in higher scores of PC1 and lower of PC2. Three species of the Linifolia clade were almost completely differentiated, which supports the previous taxonomic observations. *Iberodes linifolia* and *I. littoralis* occupied the two extremes of variation ranging from higher length of vegetative characters, higher nutlet and nutlet teeth sizes for *I. linifolia* to higher length and density of nutlet trichomes and presence of bracts for *I. littoralis* (Fig. 3a). *Iberodes kuzinskyana* occupied an intermediate position between *I. littoralis* and *I. linifolia*, overlapping very slightly with the latter. The two subspecies of *I. littoralis* were intermingled at the highest scores of PC1 and the lowest of PC2 (Fig. 3a).

### Climatic niche differentiation and distribution modelling

The two main principal components of the 6 climatic variables explained 73.59% (PC1=48.86% and PC2=24.73%) of the climatic variability in the study region. The variables that contributed most to PC1 were: bio1 (annual mean temperature), bio8 (mean temperature of wettest quarter), bio12 (annual precipitation), and bio14 (precipitation of the driest month). Bio3 (isothermality) and bio4 (temperature seasonality) were the two variables most related to PC2. The visualization of the E-space showed little overlap among taxa (Fig. 3b). The geographically widespread *Iberodes linifolia* showed the widest niche, whereas the more narrowly distributed coastal species are more restricted in climatic niche space (Fig. 3b). Among coastal species, *I. kuzinskyana* differs from *I. littoralis* in PC1 (less precipitation and higher temperatures for *I. kuzinskyana*) and overlaps with *I. littoralis* ssp*. gallaecica* but not with *I. littoralis* ssp*. littoralis* in PC2 (higher isothermality in *I. kuzinskyana* and *I. littoralis* ssp*. gallaecica*). The two subspecies of *I. littoralis* have well-differentiated niches particularly along PC2. *Iberodes kuzinskyana* occupies an intermediate position between *I. linifolia* and *I. littoralis* subsp. *gallaecica*. Thus, *I. kuzinskyana* marginally overlaps *I. linifolia* in those parts of the niche of *I. linifolia* with higher isothermality, lower temperature and higher precipitation. Indeed, minimum niche overlap was inferred between both subspecies of *I. littoralis* and *I. linifolia* due to lower temperature and higher precipitation for coastal taxa. Consequently, although tests of similarity indicated that observed values of niche overlap are not significantly lower than random overlap, all pairwise equivalency tests significantly rejected the equivalency of niches and this is supported by the low values of D obtained (Table 2).

**Table 2.** Pairwise statistical test for comparison of ecological niche overlap between the different taxa within the Linifolia clade. The statistical significance is represented by p-values (Warren *et al.*, 2008). Asterisk (*) indicate significant values for *p*-values < 0.01.

| Taxa comparison | Schoener's D | Niche equivalency p-value | Niche similarity p-value |
| --- | --- | --- | --- |
| *I. linifolia* vs *I. kuzinskyana* | 0.026 | 0.0099* | 0.94059 |
| *I. linifolia* vs *I. littoralis* ssp. *gallaecica* | 0.003 | 0.003 | 0.60396 |
| *I. linifolia* vs *I. littoralis* ssp. *littoralis* | 0.004 | 0.0099* | 0.69307 |
| *I. kuzinskyana* vs *I. littoralis* ssp. *gallaecica* | 0.000 | 0.0099* | 0.64356 |
| *I. kuzinskyana* vs *I. littoralis* ssp. *littoralis* | 0.000 | 0.0099* | 0.71287 |
| *I. littoralis* ssp. *gallaecica* vs *I. littoralis* ssp. *littoralis* | 0.000 | 0.0099* | 0.67327 |

The potential ranges for the species of the Linifolia clade in the present are broadly congruent with known geographic distributions, with overlapping distributions of *I. linifolia* and *I. kuzinskyana* (Fig. 4). In contrast, a contraction of distribution ranges was inferred for the LIG, including the disappearance of the potential range of *I. littoralis* ssp. *gallaecica* (Fig. 4), but maintaining the overlap and absence of physical barriers between *I. linifolia* and *I. kuzinskyana*. Projections to the LGM showed a range similar to the present one for *I. linifolia* and expanded ranges for both *I. kuzinskyana* and *I. littoralis* ssp. *gallaecica*. The range of *I. littoralis* ssp. *gallaecica* is inferred to have occupied most of the current range of both subspecies in the LGM, before subsp. *littoralis* differentiated from subsp. *gallaecica*.

**Fig. 4.**
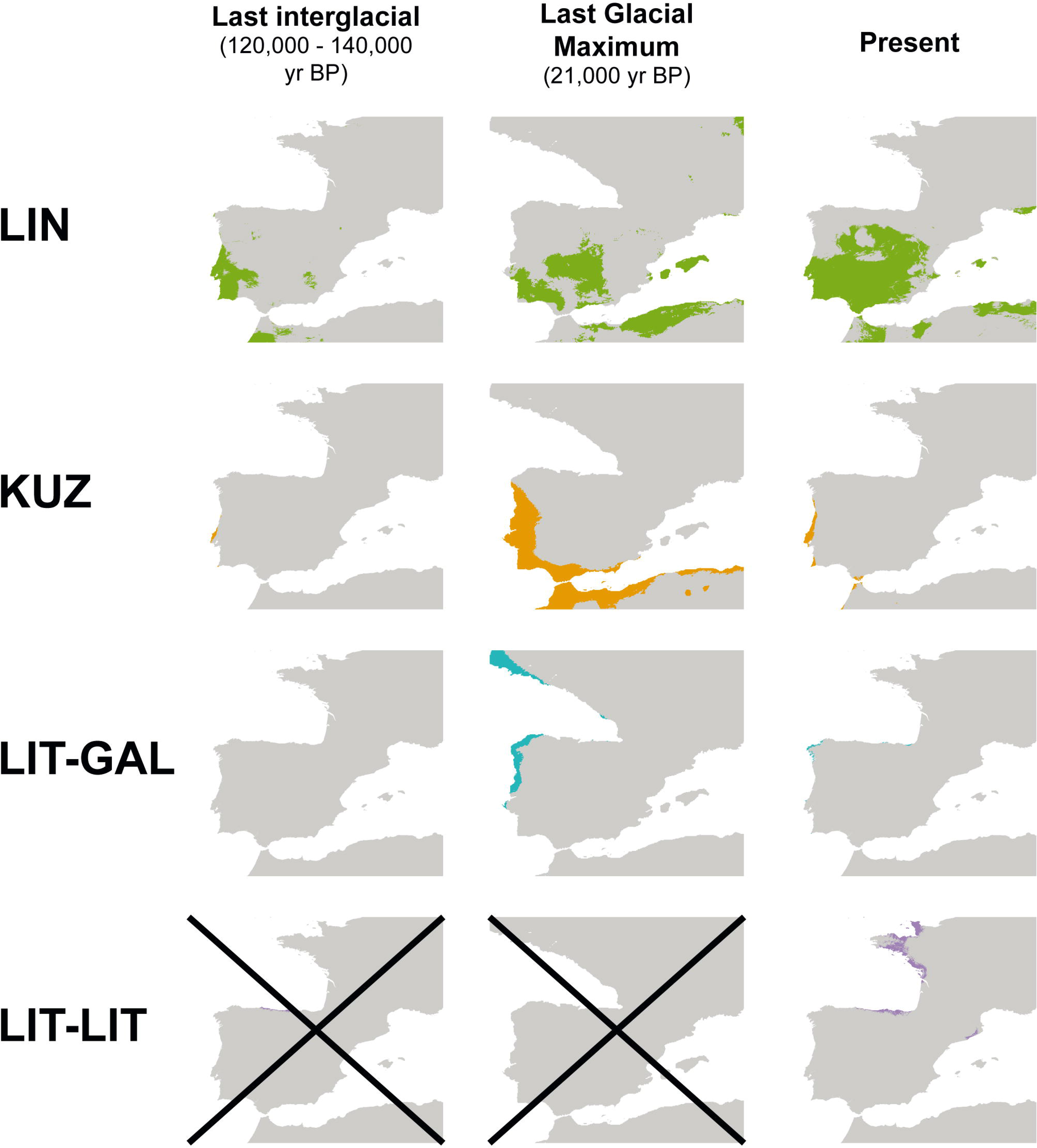
Species distribution modelling obtained through the entropy algorithm, as implemented in Maxent for the four taxa of the Linifolia clade. Codes for each taxa are: LIN for *I. linifolia*, KUZ for *I. kuzinskyana*, and LIT for *I. littoralis* (LIT-GAL for *I. littoralis* ssp. *gallaecica*, LIT-LIT for *I. littoralis* ssp. *littoralis*). Analyses are based on WorldClim and projected to past conditions: last interglacial (LIG, c. 120–140 kya) and last glacial maximum (LGM, c. 21 kya). The presence/absence was estimated using the maximum training sensitivity plus specificity logistic threshold. The projection to the past is only presented according to the inferred times of divergence of each taxa.

### Ploidy level estimation

Mean peaks of fluorescence, CV values, and genome sizes obtained for each species are summarized in Fig. S4 and Table S5. Fluorescence peaks were similar among all *Iberodes* species, ranging from 45.2 to 62.6 nm for G1. No evidence of duplicated DNA content was observed, suggesting no ploidy differences among taxa or populations. Likewise, similar genome sizes were obtained for the five species ranging from 1.788 to 1.980 pg. *Iberodes kuzinskyana* and *I. littoralis* ssp. *littoralis* exhibited the largest and the smallest genome sizes in the genus, respectively. The five species are similar in genome size to the *S. lycopersicum* standard (1.88-2.07 pg, Grandillo *et al.*, 2011).

## DISCUSSION

Phylogenetic relationships among *Iberodes* species are robustly inferred by RAD-sequencing, which demonstrated the monophyly of populations except for *I. linifolia*. Two early-diverging monophyletic species (*I. brassicifolia* and *I. commutata*) contrast with paraphyly of three species in the Linifolia clade (Fig. 1). The paraphyly of *I. linifolia* with respect to *I. kuzinskyana* and *I. littoralis* suggests a progenitor-derivative relationship between the species, with the widespread species giving rise to the coastal endemics in the course of a northward colonization from the coast of Portugal to the west coast of France. Our results provide multiple lines of evidence that support parapatric divergence of the coastland lineage (*I. kuzinskyana*-*I. littoralis* clade): (1) lack of any physical barrier to gene flow between inland and coastland in the present or the past; (2) low level of contemporary gene flow combined with evidence of historical introgression, supporting budding differentiation rather secondary contact; (3) recent niche differentiation from inland habitats with higher temperature seasonality to coastland ones with higher precipitation; (4) paraphyly of *I. linifolia*, which can be interpreted as the extant inland ancestor of the coastland lineage; and (5) incipient morphological differentiation between *I. linifolia* and the coastal lineage. Subsequently, differentiation in allopatry of the two coastal species during the mid-Pleistocene led to the modern-day species *I. kuzinskyana* and *I. littoralis*, most likely mediated by geographic isolation and niche differentiation. Thus ecological and geographic differentiation contributed to both parapatric and allopatric speciation in the clade.

### A predominant paraphyletic pattern during *Iberodes* evolution

Our RAD-seq analyses reveal a pattern of predominant paraphyly within the recently diverged Linifolia clade (Fig. 2). While the populations of *I. brassicifolia* and those of *I. commutata* form monophyletic groups, paraphyly is inferred for *I. linifolia* populations with respect to the two coastal species. While apparent paraphyly of species may be a consequence of introgression dragging populations toward the species to which they have introgressed (e.g., Eaton *et al.*, 2015), our analyses of historical introgression using *D*-statistics do not conclusively support a role of introgressive hybridization in shaping the paraphyletic topology (Methods S1), and genetic structure shows no evidence of recent admixture involving *I. linifolia* (Fig. 3c). This suggests that *Iberodes* contains some truly paraphyletic species, supporting a progenitor-derivative or “budding” speciation history. Despite the tendency in taxonomy to only consider monophyletic species, paraphyletic ones have been reported as particularly common in groups showing recent local speciation, i.e. groups composed of a ‘mother’ species that coexists in time and space with more restricted derivative species (Rajakaruna, 2018).

### Recent ecological speciation in parapatric *Iberodes*

Although parapatry has been argued to be common in plants (Rieseberg & Brouillet, 1994), few empirical examples are found on the literature (Table 3). Our study provides strong evidence for parapatry through a process of progenitor-derivative (or budding) speciation between *I. linifolia* and the ancestor of *I*. *kuzinskyana* and *I. littoralis*. Indeed, the projection of distribution models to past climatic periods during the Quaternary revealed a similar or greater degree of geographic overlap between the potential ranges of *I. linifolia* and *I. kuzinskyana* (Fig. 4), which makes allopatry unlikely (Zheng *et al.*, 2017). Peripheral populations of the widespread *I. linifolia* close to the coast of Portugal are the most closely related potential living progenitor of the coastal species (*I. kuzinskyana* and *I. littoralis*; Fig. 2), which seem to have differentiated in an adjacent area with no apparent physical barrier. Paraphyly similarly supports the hypothesis of budding (progenitor-derivative) speciation (Stuessy & Hörandl, 2013). In addition, the historical introgression inferred among *I. linifolia* and coastland species (Methods S1), does not conclusively underlie the paraphyletic topology. Moreover, our results of AMOVA and DAPC pointed to lack of recent gene flow between the nearby species (*I. kuzinskyana* and *I. linifolia*). This pattern supports strong isolation rather than secondary contacts as suggested in other cases of ecological differentiation (e.g. *Mimulus*, Stankowski *et al.*, 2015; *Stauracanthus*, Chozas *et al.*, 2017; Table 3). Nevertheless, a deeper population genetics study focused on the adjacent populations of Portugal is necessary to obtain more detailed estimates of gene flow.

**Table 3.** Examples of parapatric speciation in plants.

| Genus | Family | Event suggested | Reference |
| --- | --- | --- | --- |
| *Ephedra* | Ephedraceae | Ecological divergence mediated by bioclimatic niche | Yin *et al.* (2016) |
| *Fosterella* | Bromeliaceae | Ecological divergence mediated by polyploidization and | Paule *et al.* (2017) |
| *Helianthus* | Asteraceae | Ecological divergence mediate by bioclimatic niche | Andrew & Rieseberg (2013) |
| *Mimulus* | Phrymaceae | Ecological divergence mediated by pollinator shift | Sobel & Streisfeld (2015), Stankowski *et al.* (2015) |
| *Miscanthus* | Poaceae | Ecological divergence mediated by bioclimatic | Huang *et al.* (2014) |

|  |  | niche |  |
| --- | --- | --- | --- |
| *Oryza* | Poaceae | Ecological divergence mediated by bioclimatic niche | Zheng & Ge (2010) |
| *Populus* | Salicaceae | Ecological divergence mediated by bioclimatic niche | Zheng *et al.* (2017) |
| *Roscoea* | Zingiberaceae | Ecological divergence mediated by bioclimatic niche | Zhao *et al.* (2016) |
| *Senecio* | Asteraceae | Ecological divergence mediated by ecological niche | Richards & Ortiz-Barrientos (2016) |
| *Stauracanthus* | Fabaceae | Ecological divergence mediated by bioclimatic niche | Chozas *et al.* (2017) |
| *Columnea* | Gesneriaceae | Ecological divergence mediated by bioclimatic niche and pollinator shift | Schulte *et al.* (2015) |

In the earliest stages of parapatric speciation mediated by ecological differentiation, reproductive barriers are expected to evolve in response to selection towards locally adapted, divergent ecotypes (Sakaguchi *et al.*, 2019). The niche identity tests point to a role of ecologically-based selection in the speciation of the coastal lineage from *I. linifolia.* The coastal lineage is distributed in niches with less temperature seasonality and higher precipitation means than *I. linifolia* (Fig. 3b). Interestingly, although niche overlap is close to zero, similarity tests indicated a certain degree of similarity within the E-space, which points to recent divergence and suggests ongoing differentiation or some degree of niche conservatism. Besides, edaphic factors such as the specialization in coastal dune substrates may also have contributed to ecological differentiation, although this is a fine-scale soil variable that has not been mapped at high enough resolution to include meaningfully in niche models. The differentiation of coastal lineages (*I. kuzinskyana* and *I. littoralis*) from inland populations of *I. linifolia* seems to have started in western Iberian Peninsula during the Mid Pliocene (3.85-3.60 myr, Fig. 2), when increasing seasonality led to the origin of Mediterranean climate with its summer droughts (Suc, 1984). The onset of Mediterranean climate favored the differentiation and configuration of modern Mediterranean flora (Vargas *et al.*, 2018), which agrees with recent *Iberodes* speciation (Fig. 2). Posterior glacial/interglacial periods during the Late Pliocene promoted the expansion and contraction of geographic ranges and the formation of sources and sinks of genetic diversity for most Mediterranean species (Médail & Diadema, 2009). Interestingly, the overlapping area inferred for *I. linifolia* and *I. kuzinskyana* in the past is congruent with the location of glacial refugia (Estremadura and Beira litoral) proposed by Médail and Diadema (2009) and plant species previously studied as glacial refugial species (e.g. *Drosophyllum lusitanicum*: Müller & Deil, 2001; *Cistus monspeliensis*: Fernández-Mazuecos & Vargas, 2010; Coello *et al.*, unpublished; *Linaria elegans*: Fernández-Mazuecos & Vargas, 2013).

In addition, ecological differentiation in parapatric speciation involves divergent natural selection towards contrasting environments that promotes adaptation to different environments and subsequent barriers to gene flow (Schluter, 2001; Rundle & Nosil, 2005). The strength of divergent selection can modulate the degree of completeness of a speciation process (Nosil *et al.*, 2009). In our case, both the high interspecific genetic differentiation and clear morphological and ecological divergence (Fig. 3) between inland and coastal species may reflect strong divergent selection promoting rapid speciation (Pliocene-Pleistocene, Fig. 2). Indeed, our study reinforces the common trend for Mediterranean plants in which narrow endemics arise from ecological differentiation of peripheral populations of more widespread progenitor species (Papuga *et al.*, 2018). Moreover, this trend highlights the role of peripheral populations in setting the scene for diversification, thus revealing their conservation value (Debussche & Thompson, 2003).

### Allopatric differentiation along the Atlantic coast

An allopatric pattern is also found in the Linifolia clade. Given the current separation between ranges of the three coastal taxa (Fig. 1), allopatry seems to have triggered speciation. On the one hand, the widely separated ranges of *I. kuzinskyana* and *I. littoralis* subsp. *gallaecica* both in current times and in projections to the LIG (despite probable intermittent contact during the LGM) suggest either a colonization mediated by LDD followed by geographic isolation and niche shift (Fig. 3, 4) or a vicariant process mediated by an early expansion and contraction of distribution ranges. We find similar examples of recent allopatric speciation for both hypotheses: by LDD among coastal Mediterranean flora in the same period (e.g. Jakob *et al.*, 2007; Carnicero *et al.*, 2017; Herrando-Moraira *et al.*, 2017); and expansion/contraction of distribution ranges (e.g. Ortiz *et al.*, 2007; Lo Presti & Oberprieler, 2011; Fernández-Mazuecos & Vargas, 2011). On the other hand, the allopatric differentiation of the two subspecies of *I. littoralis* was likely preceded by an expansion of *I. littoralis* during the LGM, followed by geographic isolation as its range contracted and niche shifted, leading to the differentiation of *I. littoralis* subsp. *littoralis* in more recent times (Fig. 3, 4). In this case, the probable role of long distance dispersal (LDD) is further supported by the fact that the expanded area during the LGM does not seem to have formed a continuous range (Fig. 4). Indeed, unlike inland species of *Iberodes*, the three coastal taxa seem to have specializations for LDD in the form of uncinated hooks (Fig. 1) in the nutlet, related to the attachment to feathers and fur in other closely related Boraginaceae genera (Selvi *et al.*, 2011).

## CONCLUSIONS

Our integrative approach provides a comprehensive history of parapatric speciation, which has long posed a challenge to our understanding of ecological speciation. The integration of RAD-sequencing, morphometrics and climatic niche modelling provide strong evidence that progenitor-derivative speciation with gene flow, associated with ecological divergence, shapes the parapatry we observe today in Iberodes. Our work thus serves as a model for how integrative studies may serve as a powerful resource for investigating scenarios of non-allopatric processes in other groups of plants. Processes such as budding speciation bring to light the need to extend species concepts to encompass paraphyletic species as natural units of both taxonomic classification and evolutionary history: species in every sense of the word.

## ACKNOWLEDGEMENTS

We would like to thank those botanists and colleagues that help us with material sampling and particularly, Diana Íñigo, Paola Pérez, Celia Otero, David Vallecillo, Yurena Arjona, Lua López, Joaquín Ramírez, George Hinton, Javier Morente, Manuel Joao Pinto, Ruben Retuerto, Miguel Serrano, Antonio Rivas, Claude Dauge, Asli Koca, Peter Heenan, Enrique Rico, Ester Vega, Santiago Martín-Bravo, Íñigo Pulgar, and Juan Carlos Zamora. Likewise, we would also appreciate the worthy loans provided by numerous herbaria mentioned at supplementary tables. We gratefully acknowledge the work of the lab technicians of Real Jardín Botánico de Madrid, Yolanda Ruiz, Emilio Cano, Jose Gónzalez, Olga Popova, and Lucía Sastre. We also thank Lucía Checa and Miriam Pérez for the cytometry lab work. We are grateful for the many methodological comments of John McVay, Irene Villa-Machío, Yurena Arjona, Marcial Escudero, Tamara Villaverde, Javier Fernández-López, Alberto Coello, Santiago Martín-Bravo and Carmen Benítez-Benítez. Finally, we would like to thank the researchers of the Herbarium and Center for Tree Science at The Morton Arboretum (Lisle, Illinois, USA), in particular Marlene Hahn for lab assistance, and the Botany Department of the Field Museum (Chicago, Illinois, USA) for accommodating me at the Field Museum and handling the hosting during the lab stay in Chicago. These data were initially analyzed during a research stay by the first author in the Herbarium of The Morton Arboretum and The Field Museum. This study was supported by the Fundación General CSIC and Banco Santander as part of the project titled “Do all endangered species hold the same value?: origin and conservation of living fossils of flowering plants endemic to Spain.”

## AUTHOR CONTRIBUTIONS

AO, PJM, VV and PV contributed to developing the question and experimental design. AO led the material collection. AO, MFM and ALH conducted the data analysis. AO wrote the manuscript with the contribution of all authors in both interpretation and writing.

## SUPPORTING INFORMATION

Additional Supporting Information may be found online in the Supporting Information section at the end of the article.

**Fig. S1** RAxML trees of *Iberodes* using three different similarity percentages (85%, 90%, 95%) and four levels of minium coverage (m4, m12, m20, m28). Bootstrap support are indicated when is lower than 100.

**Fig. S2.** SVDquartets species tree of *Iberodes* using the eight main lineages obtained in the RAxML analyses.

**Fig. S3** Compoplot showing individual assignment probability to different species- groups of the Linofolia clade of *Iberodes*, considering the two subspecies of *I. littoralis* as one group (k=3). Codes for each taxa are: LIN for *I. linifolia*, KUZ for *I. kuzinskyana*, LIT-GAL for *I. littoralis* ssp. *gallaecica*, and LIT-LIT for *I. littoralis* ssp. *littoralis*.

**Fig. S4** Flow cytometry peaks for each of five samples used for each of the five species of *Iberodes* (including the two subspecies of *I. littoralis*. Peak of the standard *Solanum lycopersicum* is included. FL1 in the X axis indicates the signal intensity (530 nm / 28 nm; DAPI emission maximum=461). Y axis indicates the number of events found. Mean, mode, and median values of FL and percentage of variation coefficient is shown for each colored peak.

**Table S1** Data information and NCBI SRA accession numbers of all individuals sampled for RADseq.

**Table S2** Morphological characters analyzed. Values represent the mean of two individuals measured per herbarium sheet. LS: length of the stem, LL: length of the leaf, WL: wide of the leaf, PINFL: pedicel of inflorescence, LCFR: length of fruit calix, DN: diameter of the nutlet, LN: length of the nutlet, LM: length of the margin nutlet, LT: length of the margin teeth, DH: density of the hair, LH: length of trichome.

**Table S3** Data points for environmental analysis

**Table S4** Number of loci, SNPs retrieved for each of the 12 parameter combinations. Parameter abbreviations indicate: c, clustering threshold; m, minimum taxon coverage to consider a locus.

**Table S5** Genome size estimation for the five species of *Iberodes* (including the two subspecies of *I. littoralis*.

**Methods S1** Introgression analysis.

