## Supplementary material for "A snapshot of progenitor-derivative speciation in action in *Iberodes* (Boraginaceae)": Suplemental Material

Article title: A snapshot of progenitor-derivative speciation in action in *Iberodes* (Boraginaceae).

The following Supporting Information is available for this article:

**Fig. S1** RAxML trees of *Iberodes* using three different similarity percentages (85%, 90%, 95%) and four levels of minium coverage (m4, m12, m20, m28). Bootstrap support are indicated when is lower than 100.

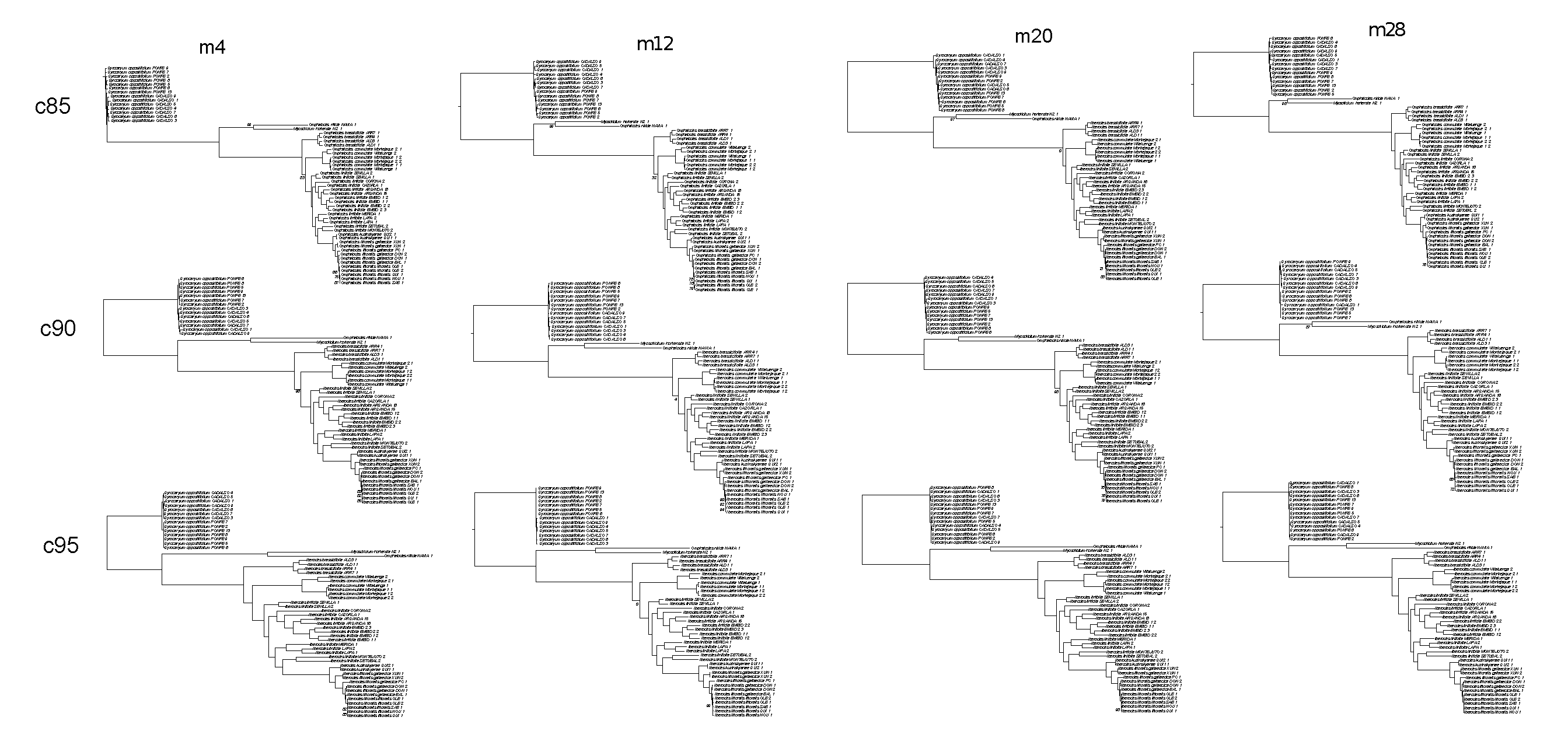
 **Fig. S2.** SVDquartets species tree of *Iberodes* using the eight main lineages obtained in the RAxML analyses.

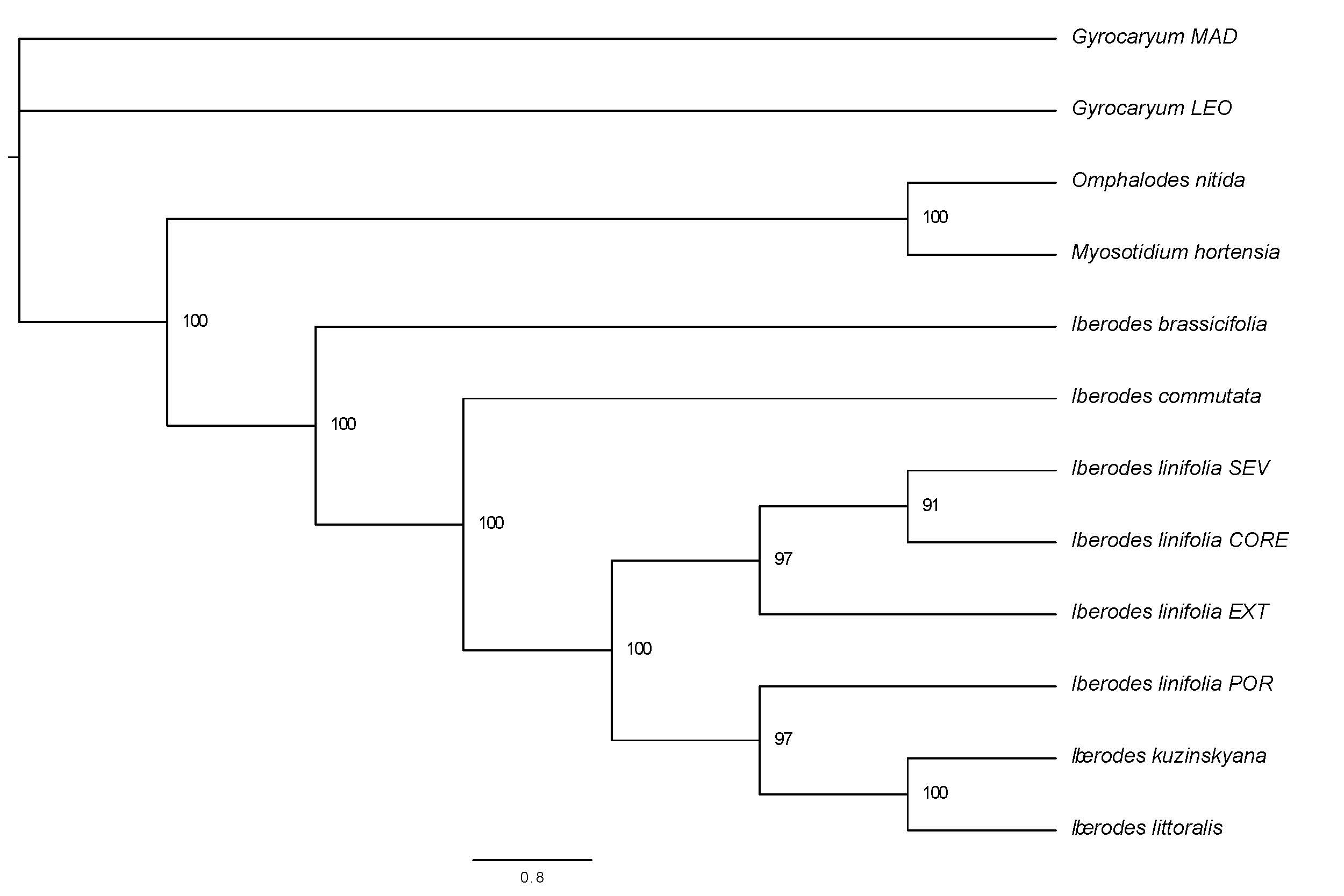

**Fig. S3** Compoplot showing individual assignment probability to different species-groups of the Linofolia clade of *Iberodes*, considering the two subspecies of *I. littoralis* as one group (k=3). Codes for each taxa are: LIN for *I. linifolia*, KUZ for *I. kuzinskyana*, LIT-GAL for *I. littoralis* ssp. *gallaecica*, and LIT-LIT for *I. littoralis* ssp. *littoralis*.

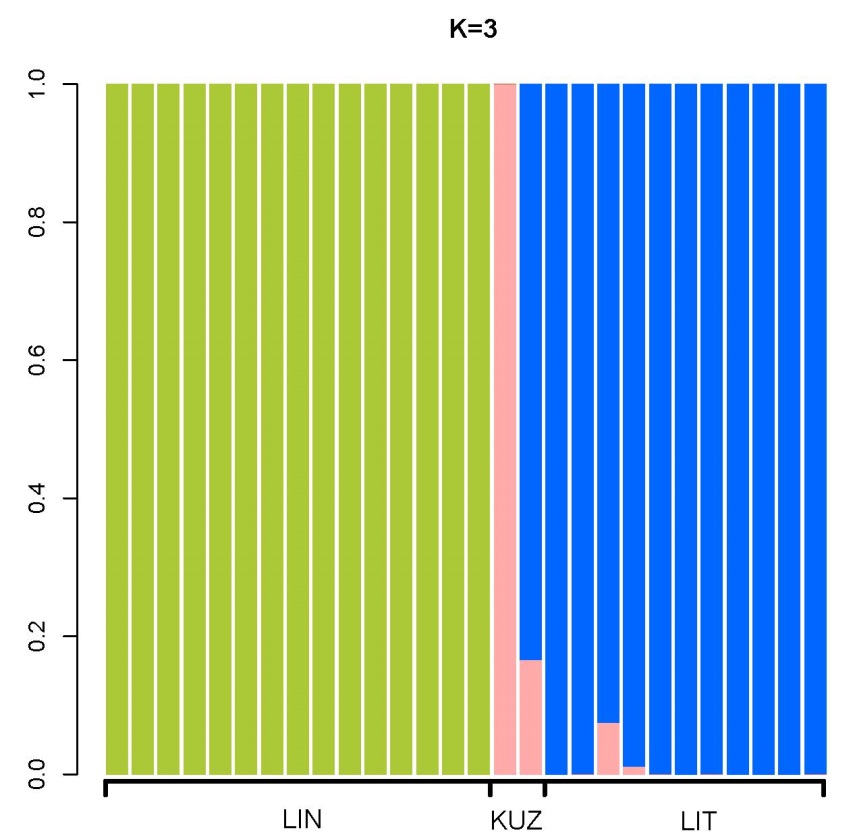

**Figure S4** Flow cytometry peaks for each of five samples used for each of the five species of *Iberodes* (including the two subspecies of *I. littoralis*. Peak of the standard *Solanum lycopersicum* is included. FL1 in the X axis indicates the signal intensity (530 nm / 28 nm; DAPI emission maximum=461). Y axis indicates the number of events found. Mean, mode, and median values of FL and percentage of variation coefficient is shown for each colored peak.

***Iberodes brassicifolia***

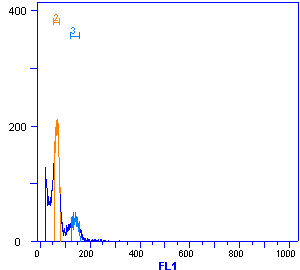

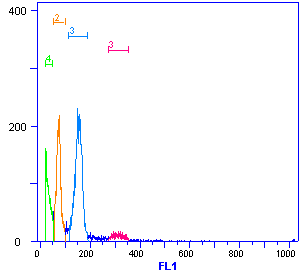

| Region | FL1 Mean | FL1 HPCV | FL1 CV | FL1 Mode | FL1 Median |
| --- | --- | --- | --- | --- | --- |
| 2 | 73.7 | 10.11% | 12.10% | 75.0 | 73.0 |
| 3 | 153.0 | 8.11% | 9.33% | 151.0 | 154.0 |
| 3 | 313.7 | 9.54% | 6.73% | 312.0 | 313.0 |
| 4 | 31.9 | 29.12% | 27.88% | 22.0 | 30.0 |

| Region | FL1 Mean | FL1 HPCV | FL1 CV | FL1 Mode | FL1 Median |
| --- | --- | --- | --- | --- | --- |
| 2 | 66.7 | 13.70% | 9.54% | 67.0 | 67.0 |
| 3 | 140.7 | 6.28% | 7.00% | 131.0 | 141.0 |

| Region | FL1 Mean | FL1 HPCV | FL1 CV | FL1 Mode | FL1 Median |
| --- | --- | --- | --- | --- | --- |
| 2 | 56.1 | 7.82% | 8.24% | 57.0 | 56.0 |
| 3 | 118.4 | 7.51% | 6.49% | 122.0 | 119.0 |
| 3 | 233.6 | 1.11% | 6.81% | 227.0 | 234.0 |

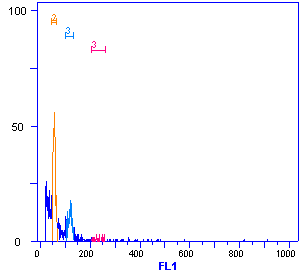

| Region | FL1 Mean | FL1 HPCV | FL1 CV | FL1 Mode | FL1 Median |
| --- | --- | --- | --- | --- | --- |
| 2 | 55.9 | 8.04% | 7.83% | 58.0 | 56.0 |
| 3 | 120.6 | 6.85% | 5.88% | 119.0 | 120.0 |
| 3 | 242.1 | 0.86% | 5.74% | 237.0 | 240.5 |

| Region | FL1 Mean | FL1 HPCV | FL1 CV | FL1 Mode | FL1 Median |
| --- | --- | --- | --- | --- | --- |
| 2 | 66.6 | 8.61% | 6.07% | 63.0 | 66.0 |
| 3 | 137.3 | 4.55% | 6.81% | 139.0 | 138.0 |

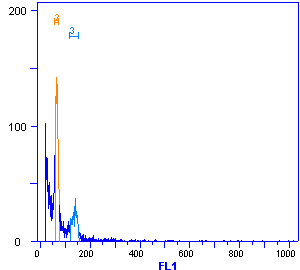

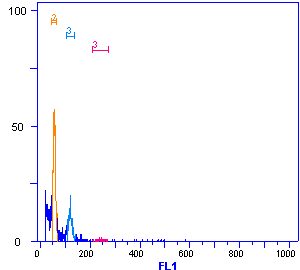

***Iberodes commutata***

| Region | FL1 Mean | FL1 HPCV | FL1 CV | FL1 Mode | FL1 Median |
| --- | --- | --- | --- | --- | --- |
| 2 | 65.5 | 8.33% | 8.40% | 66.0 | 65.0 |
| 3 | 142.4 | 1.29% | 5.90% | 133.0 | 142.0 |
| 3 | 285.1 | 2.92% | 5.62% | 284.0 | 286.0 |

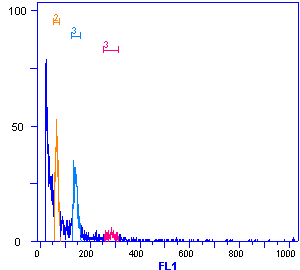

| Region | FL1 Mean | FL1 HPCV | FL1 CV | FL1 Mode | FL1 Median |
| --- | --- | --- | --- | --- | --- |
| 2 | 69.7 | 9.97% | 8.88% | 67.0 | 69.0 |
| 6 | 289.0 | 4.63% | 5.70% | 275.0 | 288.0 |
| 3 | 139.8 | 7.61% | 9.55% | 139.0 | 140.0 |

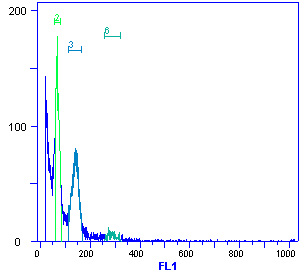

| Region | FL1 Mean | FL1 HPCV | FL1 CV | FL1 Mode | FL1 Median |
| --- | --- | --- | --- | --- | --- |
| 3 | 94.7 | 12.26% | 8.89% | 97.0 | 95.0 |
| 3 | 201.8 | 4.48% | 6.77% | 197.0 | 198.0 |
| 3 | 47.3 | 10.03% | 11.00% | 43.0 | 47.0 |

| Region | FL1 Mean | FL1 HPCV | FL1 CV | FL1 Mode | FL1 Median |
| --- | --- | --- | --- | --- | --- |
| 2 | 62.6 | 7.47% | 6.62% | 58.0 | 62.0 |
| 3 | 131.3 | 1.96% | 7.64% | 128.0 | 131.0 |
| 3 | 269.1 | 1.21% | 5.82% | 265.0 | 268.0 |
| 4 | 29.2 | 5.15% | 23.29% | 20.0 | 28.0 |

| Region | FL1 Mean | FL1 HPCV | FL1 CV | FL1 Mode | FL1 Median |
| --- | --- | --- | --- | --- | --- |
| 3 | 112.7 | 2.48% | 8.25% | 107.0 | 112.0 |
| 3 | 235.3 | 5.16% | 7.73% | 222.0 | 235.0 |

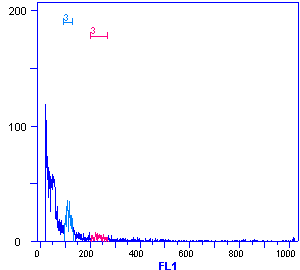

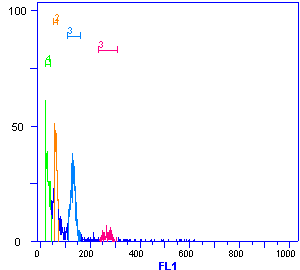

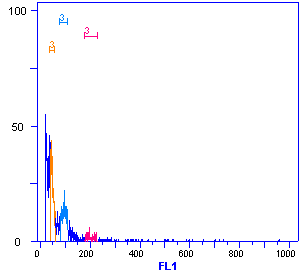

| Region | FL1 Mean | FL1 HPCV | FL1 CV | FL1 Mode | FL1 Median |
| --- | --- | --- | --- | --- | --- |
| 2 | 60.3 | 16.34% | 11.70% | 56.0 | 60.0 |
| 3 | 128.9 | 1.63% | 7.23% | 123.0 | 128.0 |
| 3 | 254.7 | 4.53% | 3.32% | 255.0 | 255.0 |

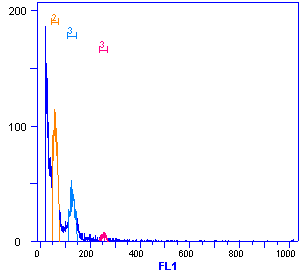
***Iberodes kuzinskyana***

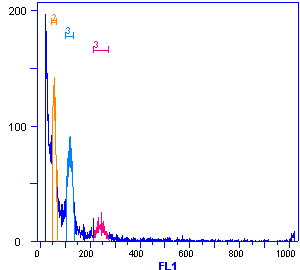

| Region | FL1 Mean | FL1 HPCV | FL1 CV | FL1 Mode | FL1 Median |
| --- | --- | --- | --- | --- | --- |
| 2 | 56.4 | 11.68% | 9.12% | 55.0 | 56.0 |
| 3 | 118.7 | 8.74% | 6.97% | 121.0 | 119.0 |
| 3 | 243.5 | 7.45% | 6.07% | 245.0 | 243.0 |

| Region | FL1 Mean | FL1 HPCV | FL1 CV | FL1 Mode | FL1 Median |
| --- | --- | --- | --- | --- | --- |
| 2 | 61.6 | 9.57% | 8.35% | 63.0 | 62.0 |
| 3 | 131.2 | 6.65% | 4.82% | 126.0 | 131.0 |
| 3 | 256.8 | 3.44% | 5.20% | 248.0 | 256.0 |

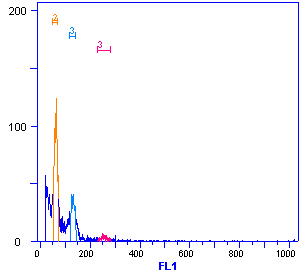

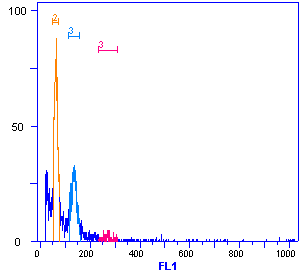

| Region | FL1 Mean | FL1 HPCV | FL1 CV | FL1 Mode | FL1 Median |
| --- | --- | --- | --- | --- | --- |
| 2 | 62.6 | 9.98% | 9.49% | 63.0 | 63.0 |
| 3 | 135.5 | 5.71% | 7.92% | 138.0 | 135.0 |
| 3 | 269.6 | 2.38% | 7.12% | 254.0 | 270.0 |

| Region | FL1 Mean | FL1 HPCV | FL1 CV | FL1 Mode | FL1 Median |
| --- | --- | --- | --- | --- | --- |
| 2 | 56.7 | 12.07% | 8.65% | 56.0 | 57.0 |
| 3 | 122.7 | 5.06% | 5.29% | 119.0 | 122.0 |
| 3 | 247.0 | 7.00% | 6.90% | 250.0 | 247.0 |

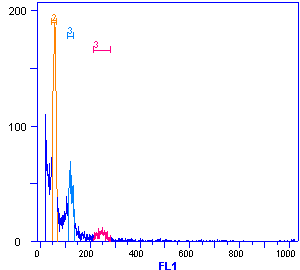

| Region | FL1 Mean | FL1 HPCV | FL1 CV | FL1 Mode | FL1 Median |
| --- | --- | --- | --- | --- | --- |
| 2 | 38.1 | 10.41% | 10.77% | 34.0 | 38.0 |
| 3 | 86.3 | 5.55% | 9.30% | 85.0 | 85.0 |
| 3 | 175.7 | 1.81% | 7.06% | 178.0 | 178.0 |

| Region | FL1 Mean | FL1 HPCV | FL1 CV | FL1 Mode | FL1 Median |
| --- | --- | --- | --- | --- | --- |
| 2 | 62.3 | 13.01% | 11.90% | 61.0 | 62.0 |
| 3 | 130.2 | 7.92% | 9.29% | 131.0 | 130.0 |
| 3 | 264.3 | 8.97% | 7.97% | 273.0 | 264.0 |
| 4 | 29.8 | 34.50% | 22.98% | 20.0 | 29.0 |

***Iberodes linifolia***

| Region | FL1 Mean | FL1 HPCV | FL1 CV | FL1 Mode | FL1 Median |
| --- | --- | --- | --- | --- | --- |
| 3 | 52.4 | 10.18% | 8.70% | 53.0 | 52.0 |
| 3 | 112.6 | 7.11% | 6.20% | 112.0 | 112.0 |
| 3 | 233.8 | 7.95% | 6.19% | 219.0 | 233.0 |

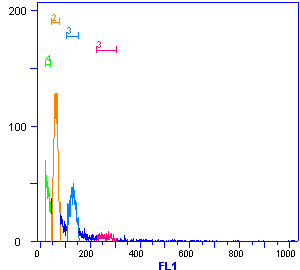

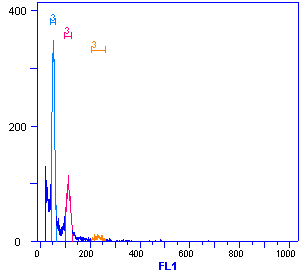

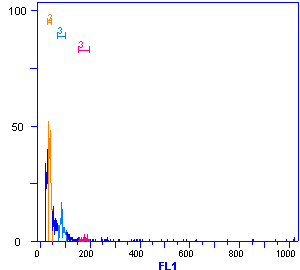

| Region | FL1 Mean | FL1 HPCV | FL1 CV | FL1 Mode | FL1 Median |
| --- | --- | --- | --- | --- | --- |
| 2 | 57.9 | 12.73% | 9.75% | 65.0 | 58.0 |
| 3 | 129.0 | 10.94% | 7.47% | 126.0 | 129.0 |
| 3 | 267.6 | 4.01% | 4.93% | 274.0 | 268.0 |

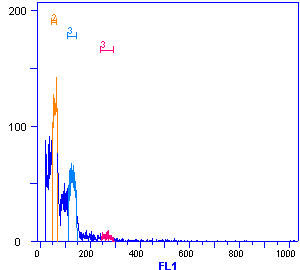

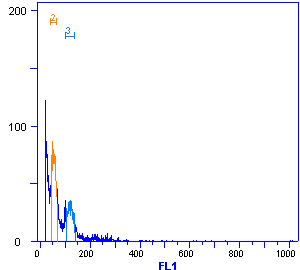

| Region | FL1 Mean | FL1 HPCV | FL1 CV | FL1 Mode | FL1 Median |
| --- | --- | --- | --- | --- | --- |
| 2 | 54.9 | 18.41% | 11.74% | 47.0 | 55.0 |
| 3 | 120.1 | 9.96% | 8.04% | 119.0 | 120.0 |

| Region | FL1 Mean | FL1 HPCV | FL1 CV | FL1 Mode | FL1 Median |
| --- | --- | --- | --- | --- | --- |
| 2 | 125.0 | 6.04% | 6.05% | 128.0 | 125.0 |
| 2 | 59.2 | 8.90% | 8.09% | 58.0 | 59.0 |
| 3 | 251.6 | 6.15% | 4.24% | 246.0 | 252.0 |

***Iberodes littoralis* ssp. *gallaecica***

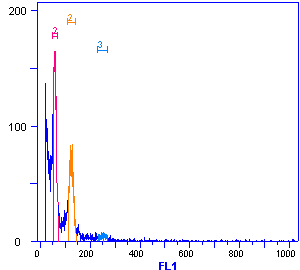

| Region | FL1 Mean | FL1 HPCV | FL1 CV | FL1 Mode | FL1 Median |
| --- | --- | --- | --- | --- | --- |
| 2 | 53.8 | 10.63% | 9.01% | 56.0 | 54.0 |
| 3 | 117.0 | 7.88% | 7.20% | 117.0 | 117.0 |
| 3 | 242.7 | 4.71% | 4.73% | 237.0 | 242.5 |

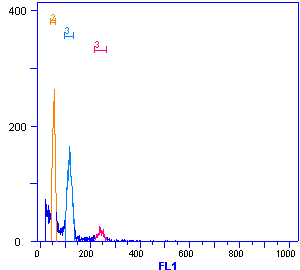

| Region | FL1 Mean | FL1 HPCV | FL1 CV | FL1 Mode | FL1 Median |
| --- | --- | --- | --- | --- | --- |
| 2 | 65.6 | 12.53% | 9.07% | 60.0 | 65.0 |
| 3 | 137.9 | 8.28% | 6.26% | 132.0 | 137.0 |
| 3 | 279.5 | 6.78% | 4.90% | 261.0 | 279.0 |

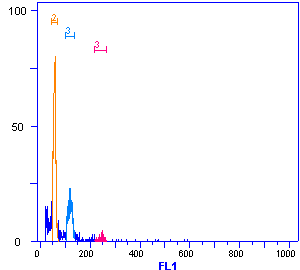

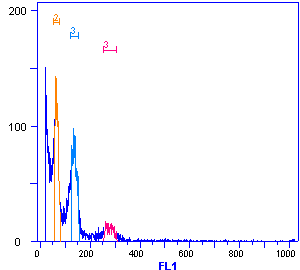

| Region | FL1 Mean | FL1 HPCV | FL1 CV | FL1 Mode | FL1 Median |
| --- | --- | --- | --- | --- | --- |
| 2 | 56.8 | 6.72% | 7.97% | 59.0 | 57.0 |
| 3 | 119.4 | 9.48% | 6.92% | 118.0 | 119.5 |
| 3 | 244.2 | 1.71% | 4.62% | 249.0 | 247.0 |

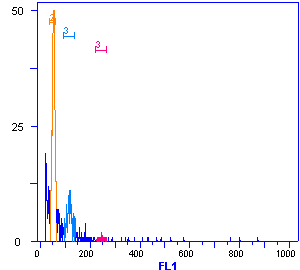

| Region | FL1 Mean | FL1 HPCV | FL1 CV | FL1 Mode | FL1 Median |
| --- | --- | --- | --- | --- | --- |
| 2 | 52.6 | 8.06% | 9.98% | 53.0 | 53.0 |
| 3 | 116.9 | 8.48% | 8.82% | 116.0 | 116.0 |
| 3 | 246.1 | 1.39% | 4.98% | 247.0 | 247.0 |

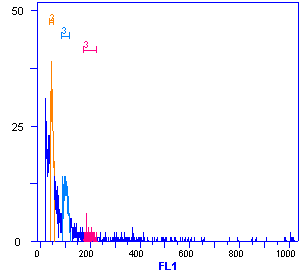
***Iberodes littoralis* ssp. *littoralis***

| Region | FL1 Mean | FL1 HPCV | FL1 CV | FL1 Mode | FL1 Median |
| --- | --- | --- | --- | --- | --- |
| 2 | 46.8 | 10.86% | 9.15% | 43.0 | 46.5 |
| 3 | 101.8 | 10.59% | 7.87% | 90.0 | 101.0 |
| 3 | 198.8 | 1.66% | 7.22% | 186.0 | 198.0 |

| Region | FL1 Mean | FL1 HPCV | FL1 CV | FL1 Mode | FL1 Median |
| --- | --- | --- | --- | --- | --- |
| 2 | 56.7 | 14.27% | 9.65% | 53.0 | 57.0 |
| 3 | 116.5 | 11.69% | 8.40% | 119.0 | 116.0 |
| 3 | 243.1 | 6.56% | 6.25% | 256.0 | 243.0 |

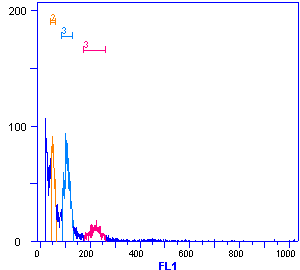

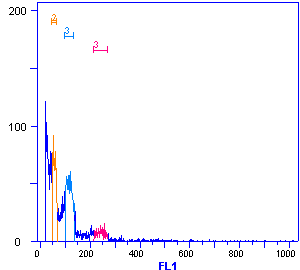

| Region | FL1 Mean | FL1 HPCV | FL1 CV | FL1 Mode | FL1 Median |
| --- | --- | --- | --- | --- | --- |
| 2 | 51.9 | 13.39% | 10.18% | 48.0 | 51.0 |
| 3 | 107.4 | 5.37% | 10.01% | 101.0 | 107.0 |
| 3 | 219.7 | 8.79% | 9.55% | 226.0 | 220.0 |

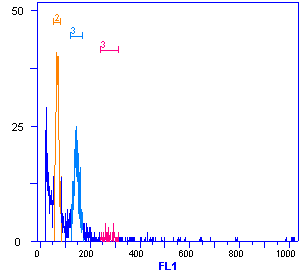

| Region | FL1 Mean | FL1 HPCV | FL1 CV | FL1 Mode | FL1 Median |
| --- | --- | --- | --- | --- | --- |
| 2 | 68.3 | 10.11% | 9.77% | 65.0 | 68.0 |
| 3 | 144.9 | 8.84% | 7.74% | 144.0 | 144.0 |
| 3 | 279.4 | 1.03% | 7.80% | 261.0 | 277.5 |

| Region | FL1 Mean | FL1 HPCV | FL1 CV | FL1 Mode | FL1 Median |
| --- | --- | --- | --- | --- | --- |
| 2 | 45.2 | 13.87% | 11.59% | 47.0 | 45.0 |
| 3 | 100.9 | 14.58% | 9.59% | 94.0 | 101.0 |
| 3 | 219.4 | 1.24% | 5.02% | 219.0 | 219.0 |

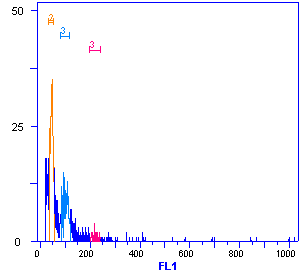

***Solanum lycopersicum***

| Region | FL1  Mean | FL1  CV | FL1  Median |
| --- | --- | --- | --- |
| 3 | 360.2 | 5.85% | 362.0 |
| 3 | 176.4 | 5.90% | 177.0 |

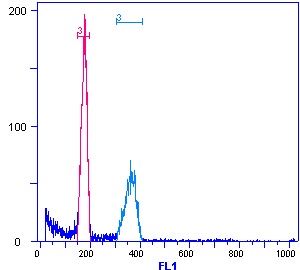

**Table S1** Data information and NCBI SRA accessions of all individuals sampled for RADseq.

| **Species** | **Population code** | **Locality** | **Coordinate X** | **Coordinate Y** | **Voucher** | **NCBI SRA** |
| --- | --- | --- | --- | --- | --- | --- |
| *Iberodes brassicifolia* | SAL 1 1 | Aldeharcipreste. Salamanca. Spain | 40.372584 | -5.887497 | AOBRA151 | Forthcoming |
| *Iberodes brassicifolia* | SAL 3 1 | Aldeharcipreste. Salamanca. Spain | 40.368197 | -5.942836 | AOBRA1529 | Forthcoming |
| *Iberodes brassicifolia* | CAC 4 1 | Arroyomolinos de la Vera. Cáceres. Spain | 40.05346 | -5.871414 | AOBRA1548 | Forthcoming |
| *Iberodes brassicifolia* | CAC 7 1 | Arroyomolinos de la Vera. Cáceres. Spain | 40.053432 | -5.87321 | AOBRA1569 | Forthcoming |
| *Iberodes commutata* | CAD 1 | Villaluenga. Cádiz. Spain | 36.6963194 | -5.3893361 | AOCOM151 (cult.) | Forthcoming |
| *Iberodes commutata* | CAD 2 | Villaluenga. Cádiz. Spain | 36.6963194 | -5.3893361 | AOCOM152 (cult.) | Forthcoming |
| *Iberodes commutata* | MAL 1 1 | Montejaque. Málaga. Spain | 36.7343472 | -5.2555778 | AOCOM156 (cult.) | Forthcoming |
| *Iberodes commutata* | MAL 1 2 | Montejaque. Málaga. Spain | 36.7343472 | -5.2555778 | AOCOM157 (cult.) | Forthcoming |
| *Iberodes commutata* | MAL 2 1 | Montejaque. Málaga. Spain | 36.7313722 | -5.2634139 | AOCOM1512 (cult.) | Forthcoming |
| *Iberodes commutata* | MAL 2 2 | Montejaque. Málaga. Spain | 36.7313722 | -5.2634139 | AOCOM1513 (cult.) | Forthcoming |
| *Iberodes kuzinskyana* | LIS 1 1 | Playa El Guincho. Cascais. Portugal | 38.740506 | -9.46984 | AOKUZ151 | Forthcoming |
| *Iberodes kuzinskyana* | LIS 2 1 | Playa El Guincho. Cascais. Portugal | 38.7401583 | -9.4699 | AOKUZ1512 | Forthcoming |
| *Iberodes linifolia* | BAD 1 | Mérida. Spain | 38.9458583 | -6.3654639 | SALA102979 | Forthcoming |
| *Iberodes linifolia* | BAD 2 | La Lapa. Sierra Alconera. Mérida. Spain | 38.4558167 | -6.5196111 | SALA125934 | Forthcoming |
| *Iberodes linifolia* | BAD 3 | La Lapa. Sierra Alconera. Mérida. Spain | 38.4558167 | -6.5196111 | SALA125934 | Forthcoming |
| *Iberodes linifolia* | BUR 2 | La Corona. Burgos. Spain | 38.4558167 | -6.5196111 | SALA150020 | Forthcoming |
| *Iberodes linifolia* | LIS 2 | Sierra do Montejuto. Portugal | 38.4558167 | -6.5196111 | SALA156157 | Forthcoming |
| *Iberodes linifolia* | ZAR 1 1 | Embid de la Ribera. Zaragoza. Spain | 40.410365 | -3.688974 | AOLIN161 | Forthcoming |
| *Iberodes linifolia* | ZAR 1 2 | Embid de la Ribera. Zaragoza. Spain | 40.410365 | -3.688974 | AOLIN162 | Forthcoming |
| *Iberodes linifolia* | ZAR 2 2 | Embid de la Ribera. Zaragoza. Spain | 40.410365 | -3.688974 | AOLIN165 | Forthcoming |
| *Iberodes linifolia* | ZAR 2 3 | Embid de la Ribera. Zaragoza. Spain | 40.410365 | -3.688974 | AOLIN166 | Forthcoming |
| *Iberodes linifolia* | SET 2 | Setúbal. Arrábida. Portugal | 38.4847 | -8.9817139 | MA730566 | Forthcoming |
| *Iberodes linifolia* | MAD 16 | San Martín de la Vega. Madrid. Spain | 40.26833 | -3.495487 | AOLIN159 | Forthcoming |
| *Iberodes linifolia* | MAD 15 | San Martín de la Vega. Madrid. Spain | 40.26833 | -3.495487 | AOLIN1519 | Forthcoming |
| *Iberodes linifolia* | JAE 1 | Cazorla. Jaén. Spain | 38.1807222 | -2.8018333 | SMB 2015 | Forthcoming |
| *Iberodes linifolia* | SEV 1 | Sevilla. Spain | 37.7381028 | -5.9845861 | I. Pulgar. 2015_1 | Forthcoming |
| *Iberodes linifolia* | SEV 2 | Sevilla. Spain | 37.7381028 | -5.9845861 | I. Pulgar. 2015_2 | Forthcoming |
| *Iberodes littoralis* subsp. *gallaecica* | DON 1 | Doniños. Coruña. Spain | 43.4992139 | -8.3162833 | AOGAL151 (cult.) | Forthcoming |
| *Iberodes littoralis* subsp. *gallaecica* | DON 2 | Doniños. Coruña. Spain | 43.4992139 | -8.3162833 | AOGAL152 (cult.) | Forthcoming |
| *Iberodes littoralis* subsp. *gallaecica* | XUN 1 | Xuño. Pontevedra. Spain | 42.631861 | -9.041028 | AOGAL158 (cult.) | Forthcoming |
| *Iberodes littoralis* subsp. *gallaecica* | XUN 2 | Xuño. Pontevedra. Spain | 42.631694 | -9.040889 | AOGAL1515 (cult.) | Forthcoming |
| *Iberodes littoralis* subsp. *gallaecica* | PC 1 | Ponteceso. Pontevedra. Spain | 43.2352722 | -8.9349361 | AOGAL1540 (cult.) | Forthcoming |
| *Iberodes littoralis* subsp. *gallaecica* | BAL 1 | Baldaio. Coruña. Spain | 43.2988611 | -8.6734028 | AOGAL1541 (cult.) | Forthcoming |
| *Iberodes littoralis* subsp*. littoralis* | POI 1 | Ile d' Oleron. La Roche. France | 45.8602056 | -1.2528722 | AOLIT151 | Forthcoming |
| *Iberodes littoralis* subsp*. littoralis* | POI 2 | Ile d' Oleron. La Roche. France | 45.9307333 | -1.3568806 | AOLIT1524 | Forthcoming |
| *Iberodes littoralis* subsp. *littoralis* | LOI 1 | Les Sables d'Olon. La Rochelle. France | 46.510131 | -1.817666 | AOLIT1568 | Forthcoming |
| *Iberodes littoralis* subsp. *littoralis* | LOI 2 | Nourmourtier. France | 46.9499694 | -2.1914444 | AOLIT1569 | Forthcoming |
| *Iberodes littoralis* subsp. *littoralis* | BRE 1 | Quiberon. France | 47.473329 | -3.090991 | AOLIT1594 | Forthcoming |
| *Gyrocaryum oppositifolium* | MAD 1 | Cadalso de los Vidrios. Madrid. Spain | 40.32043 | -4.38876 | AOGYR161 | Forthcoming |
| *Gyrocaryum oppositifolium* | MAD 3 | Cadalso de los Vidrios. Madrid. Spain | 40.32043 | -4.38876 | AOGYR163 | Forthcoming |
| *Gyrocaryum oppositifolium* | MAD 4 | Cadalso de los Vidrios. Madrid. Spain | 40.32043 | -4.38876 | AOGYR164 | Forthcoming |
| *Gyrocaryum oppositifolium* | MAD 5 | Cadalso de los Vidrios. Madrid. Spain | 40.32043 | -4.38876 | AOGYR165 | Forthcoming |
| *Gyrocaryum oppositifolium* | MAD 6 | Cadalso de los Vidrios. Madrid. Spain | 40.32043 | -4.38876 | AOGYR166 | Forthcoming |
| *Gyrocaryum oppositifolium* | MAD 7 | Cadalso de los Vidrios. Madrid. Spain | 40.32043 | -4.38876 | AOGYR167 | Forthcoming |
| *Gyrocaryum oppositifolium* | MAD 9 | Cadalso de los Vidrios. Madrid. Spain | 42.55679 | -6.54896 | AOGYR169 | Forthcoming |
| *Gyrocaryum oppositifolium* | LEO 1 | Ponferrada. León. Spain | 42.55679 | -6.54896 | AOGYR1612 | Forthcoming |
| *Gyrocaryum oppositifolium* | LEO 2 | Ponferrada. León. Spain | 42.55679 | -6.54896 | AOGYR1613 | Forthcoming |
| *Gyrocaryum oppositifolium* | LEO 5 | Ponferrada. León. Spain | 42.55679 | -6.54896 | AOGYR1616 | Forthcoming |
| *Gyrocaryum oppositifolium* | LEO 6 | Ponferrada. León. Spain | 42.55679 | -6.54896 | AOGYR1617 | Forthcoming |
| *Gyrocaryum oppositifolium* | LEO 7 | Ponferrada. León. Spain | 42.55679 | -6.54896 | AOGYR1618 | Forthcoming |
| *Gyrocaryum oppositifolium* | LEO 8 | Ponferrada. León. Spain | 42.55679 | -6.54896 | AOGYR1619 | Forthcoming |
| *Gyrocaryum oppositifolium* | LEO 9 | Ponferrada. León. Spain | 42.55679 | -6.54896 | AOGYR1620 | Forthcoming |
| *Gyrocaryum oppositifolium* | LEO 13 | Ponferrada. León. Spain | 40.372584 | -5.887497 | AOGYR1624 | Forthcoming |
| *Myosotidium hortensia* | CHA 1 | Chatham Islands. New Zealand | 40.32043 | -4.38876 | CHR609596 | Forthcoming |
| *Omphalodes nitida* | LUG 1 | Navía de Suarna. Lugo | 42.96138 | -7.009198 | AONIT151 | Forthcoming |

**Table S2** Morphological characters analyzed. Values represent the mean of two individuals measured per herbarium sheet. LS: length of the stem. LL: length of the leaf. WL: wide of the leaf. PINFL: pedicel of inflorescence. LCFR: length of fruit calix. DN: diameter of the nutlet. LN: length of the nutlet. LM: length of the margin nutlet. LT: length of the margin teeth. DH: density of the hair. LH: length of trichome.

| **Species** | **VOUCHER** | **Locality** | **LS** | **LL** | **WL** | **PINFL** | **BBR** | **LCFR** | **DN** | **LN** | **LM** | **LT** | **DH** | **LH** |
| --- | --- | --- | --- | --- | --- | --- | --- | --- | --- | --- | --- | --- | --- | --- |
| *Iberodes linifolia* | SALA108149 | Castromonte. Valladolid. Spain | 32.65 | 3.60 | 3.25 | 1.90 | 0.00 | 6.25 | 4.75 | 3.50 | 2.00 | 0.48 | 20.00 | 0.40 |
| *Iberodes linifolia* | SALA127914 | Monzón de Campos. Palencia. Spain | 28.50 | 1.80 | 2.50 | 1.25 | 0.00 | 4.75 | 5.00 | 2.25 | 1.50 | 0.38 | 7.00 | 0.25 |
| *Iberodes linifolia* | SALA10987 | Embalse de El Chorro. Málaga. Spain | 20.75 | 1.35 | 2.13 | 1.48 | 0.00 | 6.25 | 5.00 | 2.75 | 1.50 | 0.55 | 0.00 | 0.00 |
| *Iberodes linifolia* | SALA28097 | Cerro Carija. Badajoz. Spain | 19.70 | 1.83 | 2.50 | 1.00 | 0.00 | 4.25 | 4.75 | 3.00 | 1.50 | 0.48 | 0.00 | 0.00 |
| *Iberodes linifolia* | SALA39088 | Villaseca. Segovia. Spain | 22.40 | 1.75 | 1.75 | 0.90 | 0.00 | 5.00 | 3.75 | 3.00 | 1.25 | 0.58 | 0.00 | 0.00 |
| *Iberodes linifolia* | SALA57165 | Herrera de Alcántara. Cáceres. Spain | 72.50 | 3.40 | 2.75 | 3.00 | 0.00 | 6.75 | 4.50 | 3.75 | 1.50 | 0.38 | 0.00 | 0.00 |
| *Iberodes linifolia* | SALA13227 | Valdecañas de Tajo. Cáceres. Spain | 40.25 | 3.25 | 2.50 | 3.10 | 0.00 | 5.25 | 3.63 | 3.50 | 1.38 | 0.45 | 11.00 | 0.18 |
| *Iberodes linifolia* | SALA21718 | Urueña. Valladolid. Spain | 28.00 | 1.60 | 1.75 | 1.30 | 0.00 | 4.00 | 3.75 | 2.25 | 0.75 | 0.53 | 6.50 | 0.20 |
| *Iberodes linifolia* | SALA25824 | Villagarcía de Campos. Valladolid. Spain | 19.75 | 1.60 | 2.25 | 0.85 | 0.00 | 4.75 | 4.25 | 2.75 | 1.00 | 0.55 | 14.00 | 0.25 |
| *Iberodes linifolia* | SALA25869 | Urueña. Valladolid. Spain | 33.50 | 4.65 | 9.05 | 1.25 | 0.00 | 6.75 | 5.25 | 3.50 | 1.25 | 0.45 | 4.00 | 0.20 |
| *Iberodes linifolia* | SALAF4979 | Cerro Carija. Badajoz. Spain | 23.75 | 1.90 | 2.25 | 0.80 | 0.00 | 4.50 | 4.75 | 3.75 | 1.50 | 0.48 | 1.00 | 0.05 |
| *Iberodes linifolia* | SALA5283 | Puente Genil. Córdoba. Spain | 16.20 | 1.30 | 3.00 | 1.25 | 0.00 | 5.50 | 5.00 | 3.25 | 1.25 | 0.38 | 2.50 | 0.15 |
| *Iberodes linifolia* | SALA136350 | Valdajos. Toledo. Spain | 27.25 | 1.75 | 2.00 | 1.60 | 0.00 | 4.25 | 3.75 | 2.75 | 0.75 | 0.50 | 4.00 | 0.40 |
| *Iberodes linifolia* | SALA87421 | Puertollano. Ciudad Real. Spain | 30.00 | 1.75 | 2.75 | 1.08 | 0.00 | 3.25 | 3.75 | 2.25 | 1.00 | 0.38 | 0.00 | 0.00 |
| *Iberodes linifolia* | SALA26801 | Serreta de Santa Ana. Cáceres. Spain | 23.15 | 0.75 | 1.50 | 1.10 | 0.00 | 4.00 | 4.50 | 1.75 | 1.25 | 0.55 | 0.00 | 0.00 |
| *Iberodes linifolia* | SALA14448 | Ciudad Rodrigo. Salamanca. Spain | 25.00 | 1.28 | 1.75 | 1.25 | 0.00 | 4.75 | 3.35 | 2.00 | 0.75 | 0.40 | 5.00 | 0.20 |
| *Iberodes linifolia* | SALAF16476 | Finca de la Alberca. Cáceres. Spain | 29.50 | 1.35 | 2.25 | 1.30 | 0.00 | 4.50 | 3.55 | 1.75 | 0.70 | 0.40 | 0.50 | 0.05 |
| *Iberodes linifolia* | SALA21969 | Santa Espina. Valladolid. Spain | 25.25 | 1.48 | 2.50 | 1.40 | 0.00 | 4.25 | 3.75 | 2.40 | 1.10 | 0.55 | 13.00 | 0.25 |
| *Iberodes linifolia* | LEB103471 | Dueñas. Palencia. Spain | 31.75 | 1.50 | 2.50 | 0.95 | 0.00 | 4.00 | 3.88 | 3.50 | 1.50 | 0.48 | 7.00 | 0.28 |
| *Iberodes linifolia* | SALAF2922 | Serreta de Santa Ana. Cáceres. Spain | 29.00 | 1.23 | 2.50 | 1.58 | 0.00 | 4.50 | 4.13 | 3.00 | 1.38 | 0.75 | 0.50 | 0.10 |
| *Iberodes linifolia* | SALA156157 | Caldas de Monchique. Beja. Portugal | 6.50 | 1.23 | 1.75 | 0.80 | 0.00 | 3.50 | 2.25 | 1.35 | 0.50 | 0.28 | 0.00 | 0.00 |
| *Iberodes linifolia* | SALA156158 | Ribatejo. Lisboa. Portugal | 11.00 | 1.10 | 2.50 | 0.73 | 0.00 | 4.50 | 3.25 | 2.25 | 1.25 | 0.30 | 11.50 | 0.28 |
| *Iberodes linifolia* | SALA136351 | Sagres. Faro. Portugal | 5.35 | 1.10 | 1.75 | 0.35 | 0.00 | 3.00 | 3.25 | 1.50 | 0.75 | 0.35 | 0.00 | 0.00 |
| *Iberodes linifolia* | LISU158463 | Serra da Arrabida. Setubal. Portugal | 23.25 | 2.50 | 4.25 | 1.05 | 0.00 | 5.00 | 3.75 | 3.00 | 1.25 | 0.50 | 7.00 | 0.25 |
| *Iberodes linifolia* | LISUP30800 | Caldas de Monchique. Beja. Portugal | 35.00 | 4.00 | 9.00 | 1.13 | 0.00 | 4.25 | 3.95 | 2.25 | 1.25 | 0.48 | 0.00 | 0.00 |
| *Iberodes linifolia* | LISUP30794 | Serra de Monsanto. Lisboa. Portugal | 20.50 | 1.90 | 4.25 | 1.15 | 0.00 | 6.50 | 4.50 | 3.00 | 1.15 | 0.28 | 5.50 | 0.15 |
| *Iberodes linifolia* | LISUP30802 | Queluz. Lisboa. Portugal | 30.25 | 3.30 | 6.50 | 1.40 | 1.00 | 6.03 | 3.50 | 1.80 | 1.15 | 0.48 | 6.00 | 0.25 |
| *Iberodes linifolia* | LISU256043 | Serra de Alvaiazere. Leiria. Portugal | 10.75 | 1.15 | 2.25 | 0.63 | 0.00 | 3.00 | 3.00 | 1.50 | 1.00 | 0.38 | 0.00 | 0.00 |
| *Iberodes linifolia* | ALGU304 | Lagoa. Faro. Portugal | 12.25 | 1.20 | 2.55 | 0.80 | 0.00 | 3.75 | 4.25 | 2.00 | 0.75 | 0.35 | 3.00 | 0.05 |
| *Iberodes linifolia* | SEV58957 | Carratraca. Málaga. Spain | 25.75 | 2.25 | 2.25 | 1.50 | 0.00 | 4.25 | 4.00 | 2.50 | 0.75 | 0.60 | 0.00 | 0.00 |
| *Iberodes linifolia* | SEV129071 | Hinojos. Huelva. Spain | 29.00 | 3.95 | 3.00 | 0.75 | 0.00 | 5.00 | 4.50 | 3.25 | 1.00 | 0.50 | 6.50 | 0.33 |
| *Iberodes linifolia* | SEV58939 | San José del Valle. Cádiz. Spain | 21.25 | 1.60 | 3.00 | 1.30 | 0.00 | 4.25 | 4.50 | 3.00 | 1.00 | 0.45 | 7.00 | 0.45 |
| *Iberodes linifolia* | SEV81038 | Sierra de Rute. Córdoba. Spain | 23.25 | 1.60 | 2.00 | 8.25 | 0.00 | 4.50 | 4.50 | 2.75 | 1.25 | 0.60 | 6.00 | 0.05 |
| *Iberodes linifolia* | SEV58958 | La Lentejuela. Sevilla. Spain | 31.50 | 2.90 | 4.75 | 1.15 | 0.00 | 7.00 | 5.00 | 3.50 | 1.75 | 0.40 | 3.00 | 0.15 |
| *Iberodes linifolia* | SEV125267 | Caños de Meca. Cádiz. Spain | 35.25 | 2.65 | 4.00 | 0.80 | 0.00 | 5.50 | 4.50 | 2.88 | 0.88 | 0.50 | 6.50 | 0.50 |
| *Iberodes linifolia* | SEV105379 | Morón de la Frontera. Sevilla. Spain | 35.15 | 1.73 | 2.88 | 1.30 | 0.00 | 4.50 | 4.00 | 2.63 | 1.38 | 0.50 | 5.00 | 0.35 |
| *Iberodes linifolia* | SEV256122 | Puente Genil. Córdoba. Spain | 37.75 | 2.50 | 2.50 | 1.08 | 0.00 | 5.25 | 5.25 | 3.25 | 1.50 | 0.55 | 8.00 | 0.35 |
| *Iberodes linifolia* | SEV228092 | Pico Esparteros. Sevilla. Spain | 33.50 | 3.50 | 6.00 | 1.20 | 0.00 | 6.00 | 4.75 | 3.50 | 1.38 | 0.50 | 2.00 | 0.28 |
| *Iberodes linifolia* | SEV88021 | El Burgo. Málaga. Spain | 16.85 | 3.00 | 5.00 | 0.83 | 0.00 | 5.00 | 4.50 | 3.50 | 1.00 | 0.50 | 7.00 | 0.50 |
| *Iberodes kuzinskyana* | LISU147336 | Praia do Cavalho. Lisboa. Portugal | 6.00 | 1.23 | 2.75 | 1.00 | 1.00 | 5.50 | 3.75 | 2.25 | 1.25 | 0.30 | 7.50 | 0.53 |
| *Iberodes kuzinskyana* | LISUP45600 | Praia da Ursa. Lisboa. Spain | 6.25 | 2.05 | 4.50 | 0.95 | 1.00 | 3.50 | 3.63 | 2.25 | 1.63 | 0.65 | 6.50 | 0.35 |
| *Iberodes kuzinskyana* | LISUP30805 | Cabo da Roca. Lisboa. Portugal | 6.50 | 1.75 | 4.25 | 1.30 | 1.00 | 4.00 | 4.00 | 2.40 | 1.15 | 0.45 | 6.50 | 0.25 |
| *Iberodes kuzinskyana* | LISUP30807 | Cabo da Roca. Lisboa. Portugal | 5.90 | 1.78 | 3.00 | 1.15 | 1.00 | 4.25 | 4.50 | 2.50 | 1.40 | 0.45 | 3.00 | 0.45 |
| *Iberodes kuzinskyana* | LISUP56554 | São João de Estoril. Lisboa. Portugal | 9.75 | 1.38 | 6.75 | 1.08 | 1.00 | 4.50 | 4.25 | 2.15 | 1.00 | 0.40 | 5.00 | 0.48 |
| *Iberodes kuzinskyana* | LISUP30811 | São João de Estoril. Lisboa. Portugal | 8.55 | 1.68 | 6.50 | 1.25 | 1.00 | 4.25 | 4.75 | 2.25 | 1.00 | 0.25 | 5.00 | 0.25 |
| *Iberodes kuzinskyana* | LISU151707 | Praia do Guincho. Lisboa. Portugal | 2.15 | 1.35 | 3.25 | 0.65 | 1.00 | 4.00 | 4.75 | 2.00 | 1.00 | 0.38 | 4.50 | 0.33 |
| *Iberodes littoralis* subsp. *littoralis* | MA471282 | Quiberon. Bretagne. France | 4.50 | 2.15 | 3.50 | 1.15 | 1.00 | 3.25 | 2.50 | 2.75 | 1.00 | 0.40 | 7.00 | 0.23 |
| *Iberodes littoralis* subsp. *littoralis* | MA94699 | Quiberon. Bretagne. France | 4.75 | 2.10 | 2.10 | 0.75 | 1.00 | 4.00 | 2.50 | 3.60 | 0.75 | 0.18 | 8.50 | 0.50 |
| *Iberodes littoralis* subsp. *gallaecica* | MA387710 | Baldaio. Coruña. Spain | 2.60 | 1.35 | 2.88 | 0.80 | 1.00 | 3.50 | 2.75 | 4.25 | 0.88 | 0.15 | 12.50 | 0.40 |
| *Iberodes littoralis* subsp. *gallaecica* | MA730567 | Ponteceso. Coruña. Spain | 7.05 | 1.90 | 3.13 | 1.73 | 1.00 | 6.00 | 2.38 | 2.88 | 0.88 | 0.30 | 9.50 | 0.48 |
| *Iberodes littoralis* subsp. *gallaecica* | MA469667 | Baldaio. Coruña. Spain | 4.10 | 1.60 | 2.25 | 0.85 | 1.00 | 4.75 | 2.75 | 3.25 | 1.00 | 0.35 | 5.00 | 0.35 |
| *Iberodes littoralis* subsp. *gallaecica* | MA350088 | Praia da Pedrosa. Coruña. Spain | 6.25 | 1.68 | 5.00 | 1.03 | 1.00 | 4.25 | 2.50 | 2.88 | 1.00 | 0.23 | 10.50 | 0.48 |
| *Iberodes littoralis* subsp. *gallaecica* | MA550996 | Praia de San Xurxo. Coruña. Spain | 10.03 | 2.45 | 5.25 | 1.90 | 1.00 | 5.25 | 3.63 | 3.75 | 1.13 | 0.25 | 8.00 | 0.45 |
| *Iberodes littoralis* subsp. *gallaecica* | MA387709 | Doniños. Coruña. Spain | 6.18 | 1.33 | 2.25 | 1.03 | 1.00 | 5.50 | 2.63 | 3.13 | 0.88 | 0.25 | 7.00 | 0.40 |
| *Iberodes littoralis* subsp. *littoralis* | AOLIT15113 | Quiberon. Bretagne. France | 6.55 | 2.00 | 3.13 | 1.00 | 1.00 | 4.25 | 2.38 | 3.00 | 0.88 | 0.08 | 4.50 | 0.25 |
| *Iberodes littoralis* subsp. *littoralis* | AOLIT15114 | Nourmoutier. La Loire. France | 6.33 | 1.50 | 2.50 | 1.00 | 1.00 | 4.00 | 3.25 | 3.50 | 1.10 | 0.05 | 9.00 | 0.20 |
| *Iberodes littoralis* subsp. *littoralis* | AOLIT15115 | Île d’Oleron. Poitou-Charentes. France | 8.00 | 1.50 | 3.00 | 1.00 | 1.00 | 4.00 | 2.75 | 3.00 | 0.75 | 0.20 | 8.00 | 0.40 |
| *Iberodes littoralis* subsp. *gallaecica* | AOGAL1510 | Xuño. Coruña. Spain | 5.70 | 1.25 | 2.50 | 0.90 | 1.00 | 4.00 | 2.00 | 3.00 | 1.00 | 0.50 | 9.00 | 0.40 |

**Table S3** Data points for environmental analysis

| *Species* | X | Y |
| --- | --- | --- |
| *Iberodes kuzinskyana* | -9.498028 | 38.781009 |
| *Iberodes kuzinskyana* | -9.491894 | 38.790468 |
| *Iberodes kuzinskyana* | -9.482000 | 38.802610 |
| *Iberodes kuzinskyana* | -9.482000 | 38.712500 |
| *Iberodes kuzinskyana* | -9.482000 | 38.802610 |
| *Iberodes kuzinskyana* | -9.470623 | 38.741533 |
| *Iberodes kuzinskyana* | -9.420453 | 38.930540 |
| *Iberodes kuzinskyana* | -9.403590 | 38.893180 |
| *Iberodes kuzinskyana* | -9.402570 | 38.712950 |
| *Iberodes kuzinskyana* | -9.385946 | 38.701342 |
| *Iberodes kuzinskyana* | -8.903036 | 38.487628 |
| *Iberodes linifolia* | -9.171890 | 38.443160 |
| *Iberodes linifolia* | -9.171277 | 38.708726 |
| *Iberodes linifolia* | -9.057880 | 39.164200 |
| *Iberodes linifolia* | -8.942700 | 38.485000 |
| *Iberodes linifolia* | -8.825480 | 39.524520 |
| *Iberodes linifolia* | -8.718360 | 37.181100 |
| *Iberodes linifolia* | -8.592780 | 39.523940 |
| *Iberodes linifolia* | -8.493650 | 37.103000 |
| *Iberodes linifolia* | -8.369760 | 38.441590 |
| *Iberodes linifolia* | -8.141670 | 38.350020 |
| *Iberodes linifolia* | -8.042470 | 37.177570 |
| *Iberodes linifolia* | -7.815800 | 37.265650 |
| *Iberodes linifolia* | -7.658690 | 39.697110 |
| *Iberodes linifolia* | -7.553310 | 39.155240 |
| *Iberodes linifolia* | -7.477550 | 37.261790 |
| *Iberodes linifolia* | -7.348950 | 37.981060 |
| *Iberodes linifolia* | -7.340800 | 38.341420 |
| *Iberodes linifolia* | -7.321740 | 37.751722 |
| *Iberodes linifolia* | -7.214217 | 39.443116 |
| *Iberodes linifolia* | -7.191260 | 39.688766 |
| *Iberodes linifolia* | -7.102724 | 38.697567 |
| *Iberodes linifolia* | -7.036970 | 41.039520 |
| *Iberodes linifolia* | -7.001641 | 37.909480 |
| *Iberodes linifolia* | -6.842793 | 39.684132 |
| *Iberodes linifolia* | -6.801276 | 40.948795 |
| *Iberodes linifolia* | -6.671219 | 37.878738 |
| *Iberodes linifolia* | -6.648968 | 38.509113 |
| *Iberodes linifolia* | -6.637144 | 38.869282 |
| *Iberodes linifolia* | -6.581265 | 40.578790 |
| *Iberodes linifolia* | -6.491247 | 39.678773 |
| *Iberodes linifolia* | -6.423004 | 38.414257 |
| *Iberodes linifolia* | -6.372798 | 37.956867 |
| *Iberodes linifolia* | -6.241250 | 40.118784 |
| *Iberodes linifolia* | -6.201109 | 38.228966 |
| *Iberodes linifolia* | -6.180114 | 38.769043 |
| *Iberodes linifolia* | -6.161186 | 36.428719 |
| *Iberodes linifolia* | -6.151190 | 36.698724 |
| *Iberodes linifolia* | -5.968330 | 36.180560 |
| *Iberodes linifolia* | -5.931465 | 37.856183 |
| *Iberodes linifolia* | -5.909651 | 38.136177 |
| *Iberodes linifolia* | -5.881196 | 37.328737 |
| *Iberodes linifolia* | -5.861239 | 39.848783 |
| *Iberodes linifolia* | -5.851185 | 36.698727 |
| *Iberodes linifolia* | -5.823676 | 38.859049 |
| *Iberodes linifolia* | -5.772161 | 36.449941 |
| *Iberodes linifolia* | -5.596312 | 38.032174 |
| *Iberodes linifolia* | -5.585225 | 36.684819 |
| *Iberodes linifolia* | -5.461203 | 38.238757 |
| *Iberodes linifolia* | -5.459913 | 37.745812 |
| *Iberodes linifolia* | -5.421185 | 37.158739 |
| *Iberodes linifolia* | -5.391180 | 36.898735 |
| *Iberodes linifolia* | -5.259670 | 37.834456 |
| *Iberodes linifolia* | -5.242261 | 38.691463 |
| *Iberodes linifolia* | -5.184265 | 41.731210 |
| *Iberodes linifolia* | -5.181170 | 36.528731 |
| *Iberodes linifolia* | -5.081178 | 37.158742 |
| *Iberodes linifolia* | -4.981258 | 41.758832 |
| *Iberodes linifolia* | -4.797542 | 36.600391 |
| *Iberodes linifolia* | -4.792013 | 37.085060 |
| *Iberodes linifolia* | -4.781168 | 36.838740 |
| *Iberodes linifolia* | -4.700531 | 37.851073 |
| *Iberodes linifolia* | -4.521242 | 41.398829 |
| *Iberodes linifolia* | -4.511254 | 42.038843 |
| *Iberodes linifolia* | -4.411197 | 39.018783 |
| *Iberodes linifolia* | -4.401159 | 36.718742 |
| *Iberodes linifolia* | -4.394970 | 36.955930 |
| *Iberodes linifolia* | -4.221199 | 39.338791 |
| *Iberodes linifolia* | -4.211193 | 38.978784 |
| *Iberodes linifolia* | -4.111182 | 38.488776 |
| *Iberodes linifolia* | -4.031246 | 42.078849 |
| *Iberodes linifolia* | -3.991202 | 39.788802 |
| *Iberodes linifolia* | -3.975333 | 38.213050 |
| *Iberodes linifolia* | -3.881208 | 40.238812 |
| *Iberodes linifolia* | -3.871228 | 41.318834 |
| *Iberodes linifolia* | -3.871188 | 39.068789 |
| *Iberodes linifolia* | -3.811163 | 37.628764 |
| *Iberodes linifolia* | -3.791178 | 38.558780 |
| *Iberodes linifolia* | -3.741169 | 38.078772 |
| *Iberodes linifolia* | -3.661221 | 41.148833 |
| *Iberodes linifolia* | -3.641163 | 37.858769 |
| *Iberodes linifolia* | -3.621150 | 37.088756 |
| *Iberodes linifolia* | -3.611212 | 40.698824 |
| *Iberodes linifolia* | -3.531197 | 39.968810 |
| *Iberodes linifolia* | -3.521169 | 38.348779 |
| *Iberodes linifolia* | -3.511227 | 41.608844 |
| *Iberodes linifolia* | -3.501176 | 38.798788 |
| *Iberodes linifolia* | -3.441800 | 40.271300 |
| *Iberodes linifolia* | -3.441232 | 41.938852 |
| *Iberodes linifolia* | -3.291194 | 40.068815 |
| *Iberodes linifolia* | -3.181223 | 41.778852 |
| *Iberodes linifolia* | -3.061169 | 38.888794 |
| *Iberodes linifolia* | -3.031239 | 42.738875 |
| *Iberodes linifolia* | -3.011206 | 41.008837 |
| *Iberodes linifolia* | -2.941214 | 41.508848 |
| *Iberodes linifolia* | -2.941189 | 40.158820 |
| *Iberodes linifolia* | -2.831233 | 42.638874 |
| *Iberodes linifolia* | -2.831140 | 37.358769 |
| *Iberodes linifolia* | -2.821180 | 39.798814 |
| *Iberodes linifolia* | -2.788982 | 37.876834 |
| *Iberodes linifolia* | -2.711160 | 38.708794 |
| *Iberodes linifolia* | -2.701183 | 40.098821 |
| *Iberodes linifolia* | -2.601138 | 37.538774 |
| *Iberodes linifolia* | -2.581210 | 41.688856 |
| *Iberodes linifolia* | -2.451151 | 38.468793 |
| *Iberodes linifolia* | -2.341203 | 41.588856 |
| *Iberodes linifolia* | -2.341195 | 41.138847 |
| *Iberodes linifolia* | -2.031139 | 38.258793 |
| *Iberodes linifolia* | -2.011172 | 40.238831 |
| *Iberodes linifolia* | -1.981196 | 41.588860 |
| *Iberodes linifolia* | -1.891177 | 40.638841 |
| *Iberodes linifolia* | -1.631184 | 41.318858 |
| *Iberodes linifolia* | 0.078853 | 41.198873 |
| *Iberodes linifolia* | 4.822680 | 44.458990 |
| *Iberodes linifolia* | 5.189150 | 44.182180 |
| *Iberodes linifolia* | 5.203890 | 43.978060 |
| *Iberodes littoralis gallaecica* | -9.181364 | 43.128814 |
| *Iberodes littoralis gallaecica* | -9.061364 | 43.181000 |
| *Iberodes littoralis gallaecica* | -9.061354 | 42.768808 |
| *Iberodes littoralis gallaecica* | -9.061350 | 42.588804 |
| *Iberodes littoralis gallaecica* | -9.011347 | 42.478802 |
| *Iberodes littoralis gallaecica* | -8.941362 | 43.214000 |
| *Iberodes littoralis gallaecica* | -8.941348 | 42.588805 |
| *Iberodes littoralis gallaecica* | -8.691359 | 43.288824 |
| *Iberodes littoralis gallaecica* | -8.571357 | 43.308825 |
| *Iberodes littoralis gallaecica* | -8.321356 | 43.488832 |
| *Iberodes littoralis gallaecica* | -8.201351 | 43.358831 |
| *Iberodes littoralis litoralis* | -3.999250 | 47.708000 |
| *Iberodes littoralis litoralis* | -3.260000 | 47.364700 |
| *Iberodes littoralis litoralis* | -3.112220 | 47.598330 |
| *Iberodes littoralis litoralis* | -3.085000 | 47.468500 |
| *Iberodes littoralis litoralis* | -2.890000 | 47.339000 |
| *Iberodes littoralis litoralis* | -2.303730 | 46.692630 |
| *Iberodes littoralis litoralis* | -2.295000 | 47.021000 |
| *Iberodes littoralis litoralis* | -2.059310 | 46.884130 |
| *Iberodes littoralis litoralis* | -1.911910 | 46.710090 |
| *Iberodes littoralis litoralis* | -1.773340 | 46.625740 |
| *Iberodes littoralis litoralis* | -1.627560 | 46.451330 |
| *Iberodes littoralis litoralis* | -1.482710 | 46.238000 |
| *Iberodes littoralis litoralis* | -1.345900 | 46.191850 |
| *Iberodes littoralis litoralis* | -1.337000 | 46.012020 |
| *Iberodes littoralis litoralis* | -1.236000 | 45.926970 |
| *Iberodes littoralis litoralis* | -1.090000 | 46.078110 |

**Table S4** Number of loci. SNPs retrieved for each of the 12 parameter combinations. Parameter abreviations indicate: c. clustering threshold; m. minimum taxon coverage to consider a locus.

| **Parameter Combination** | **Loci** | **SNPs** |
| --- | --- | --- |
| c85m4 | 93420 | 478361 |
| c85m12 | 43781 | 362327 |
| c85m20 | 30902 | 288146 |
| c85m28 | 22775 | 219227 |
| c90m4 | 105125 | 456317 |
| c90m12 | 49392 | 356412 |
| c90m20 | 35070 | 288996 |
| c90m28 | 25713 | 218214 |
| c95m4 | 148890 | 437879 |
| c95m12 | 64954 | 337222 |
| c95m20 | 43691 | 267158 |
| c95m28 | 30725 | 193142 |

**Table S5** Genome size estimation for the five species of *Iberodes* (including the two subspecies of *I. littoralis*.

| **Taxon** | **2C** |
| --- | --- |
| *Iberodes brassicifolia* | 1.968 |
| *Iberodes commutata* | 1.922 |
| *Iberodes linifolia* | 1.837 |
| *Iberodes kunzinskyanae* | 1.980 |
| *Iberodes littoralis* subsp. *gallaecica* | 1.984 |
| *Iberodes littoralis* subsp. *littoralis* | 1.788 |

**Methods S1**

**Introgression analysis**

Besides genetic structure analyses that test for recent admixture of taxa it is important to test also for ancestral introgression that may be masking the actual topology of the phylogenetic reconstruction. Indeed, some lineages may look paraphyletic if there is some gene flow between them, even if they are truly reciprocally monophyletic (Eaton et al., 2015). Thus, four-taxon *D*-statistic analysis (Durand et al., 2011) was conducted through *PyRAD* v. 3.0.6 to test for ancestral introgression between lineages of *I. linifolia* and coastal taxa that together form the Linifolia clade. We used the HBS matrix of concatenated loci as the input and set 200 bootstrap replicates. *D*-statistic analysis compares the occurrence of “ABBA” versus “BABA” pattern based on a four taxon pectinate topology: [(((P1, P2), P3), O)]. Ancestral introgression either between P1 and P3 (BABA) or between P2 and P3 (ABBA) is resulted if one of those patterns is significantly more frequent than the other (Eaton et al., 2015). In our case, the outgroup taxa (O) included all individuals of *I. commutata*, that showed to be the ancestral taxa of Linifolia clade. P1 and P2 corresponded to the two coastal sister species (*I. littoralis* and *I. kuzinsyana*, respectively) and individuals of *I. linifolia* as P3, the ancestor of coastal taxa.

As a result, D-statistic analyses showed some level of introgression (both ABBA and BABA patterns) between coastal species and populations of *I. linifolia*, although this introgression did not explain the paraphyletic pattern of Linifolia clade. Out of 1980 comparisons, 701 supported ABBA and 1104 BABA and only 175 were not significant (Table 1).

**Table 1.** D-statistic tests given a four-taxon tree (((P1,P2),P3),O). Percentages of significant ABBA or BABA patterns are given for the tests between each lineage of *I. linifolia* (P3) and coastal species, *I. kuzinskyana* (P2) and *I. littoralis* (P1). Abbreviations of the three taxa are: LIN, *I. linifolia*; KUZ, *I. kuzinskyana*; LIT, *I. littoralis*. Asterisk (*) indicates the introgression pair taxa. Abbreviations for each individual of *I. linifolia* lineages are described at Appendix S1a and Fig. 2 of the main text.

| ***I.linifolia* lineages (LIN)**  (P3) | **% Sig. ABBA pattern**  LIN*KUZ (P2) | **% Sig. BABA pattern**  LIN*LIT (P1) | **TOTAL** |
| --- | --- | --- | --- |
| Portugal (LIS 2, SET 2) | 49.666 | 42.727 | 330 |
| Extremadura  (BAD 1, BAD 2, BAD 3) | 42.152 | 46.637 | 446 |
| Core  (BUR 2, JAE 1, MAD 15, MAD 16, ZAR 2 2, ZAR, 2 3, ZAR 1 3 , ZAR 1 1) | 23.229 | 70.105 | 960 |
| Sevilla  (SEV 1, SEV 2) | 57.787 | 33.607 | 244 |
| TOTAL | 35.404 | 55.758 | 1980 |
